# Hypoxic stress granules trigger immunogenic dormancy in lung cancer

**DOI:** 10.64898/2026.02.18.706626

**Authors:** Matthew G. Smith, Alexis Rebecca Ramos, Heena Panchal, N. Harun Cerkezi, Cassandra Garcia, Lynn Spruce, Hossein Fazalinia, Leonard B. Maggi, Adam W. Mailloux

## Abstract

Induction of the MHC class I antigen processing and presentation pathway (C1APP) is a critical part of the IFN-γ response necessary for effective cytotoxic immunity against tumors of epithelial origin^1,2^. Loss of this response is associated with worse disease outcomes and renders patients refractory to immunotherapies^3–6^. Without C1APP induction, tumor cells cannot optimally process and present immunopeptides from tumor-associated antigens (TAA) and neoantigens to effector cytotoxic T cells^7–9^. Here, we show that physiologic levels of hypoxia block induction of the immunoproteasome (IP) and other C1APP components in cancer cells, including human non-small cell lung cancer (NSCLC). In A549 cells, this leads to impaired presentation of more than 73% of detectable immunopeptides, including TAA and neoantigen-derived immunopeptides. This effect is independent of HIF-1α or HIF-2α signaling, protein degradation, autophagy, or stimulus type. Instead, hypoxia induces translational arrest of C1APP mRNAs prior to complete monosome loading, along with sequestration into hypoxia-associated stress granules. This phenomenon is reversible with the epitranscriptomic compound 5-azacytidine. Consistent with these findings, IP expression is excluded from hypoxic regions in most human NSCLC tumors. Together, these results link tumor hypoxia to a state of “immunogenic dormancy” and identify stress granules as a previously unrecognized mechanism of immune escape.

## Main

Effective CD8⁺ T cell–mediated tumor immunity requires that cancer cells display unique immunopeptides in the context of MHC class I^10^. These peptides classically derive from both tumor-associated antigens (TAAs), including aberrantly expressed self-proteins, or mutation-derived neoantigens generated during malignant transformation. Antigen presentation depends on the class I antigen processing and presentation pathway (C1APP), in which cytosolic proteins are degraded by the proteasome into peptides that are transported into the endoplasmic reticulum, loaded on to nascent MHC class I molecules, and trafficked to the cell surface^11,12^. Inflammatory cues, particularly IFN-γ, remodel this pathway by inducing alternative catalytic β subunits (β1i, β2i, β5i) of the immunoproteasome (IP), which more efficiently produce high-affinity MHC class I peptides than constitutive subunits (β1, β2, β5)^13,14^, as well as downstream C1APP components, thereby expanding and reshaping the immunopeptidome toward peptides that are efficiently presented to cytotoxic T cells^15^.

In epithelia and epithelial-derived tumors, basal C1APP expression is typically low for all components, rendering antigen presentation highly dependent on inducible responses to immune pressure^15^. Accordingly, tumor immune visibility often hinges on whether cancer cells can be IFN-γ-responsive at the time of T cell engagement. Consistent with this model, IFN-γ-driven C1APP is strongly associated with clinical benefit from immune checkpoint blockade across multiple cancer types, identifying C1APP inducibility as a critical determinant of effective anti-tumor immunity^1,2^.

Understanding how tumors moderate the C1APP is critical. Recent insights report reduced baseline surface MHC class I through altered intracellular trafficking and lysosomal degradation, including autophagy-dependent pathways that divert existing MHC-I away from the cell surface^16^. Hypoxia may drive this autophagy-dependent loss of baseline C1APP^17^. While these observations clarify how steady-state MHC-I abundance can be reduced under stress, they remain focused on degradation and trafficking of existing C1APP components. They do not address whether tumor cells retain the capacity to induce the C1APP in response to inflammatory cues. In epithelial tumors, where basal C1APP activity is low, any impairment of this inducible response represents a distinct and consequential limitation on CD8⁺ T cell recognition.

Here, we demonstrate that exposure to 2% O₂ (a condition commonly used to model mild hypoxia^18^) is sufficient to block IFN-γ-mediated induction of the C1APP and impair downstream immunopeptide presentation in human lung cancer cells. Lung cancer represents a particularly relevant model, as it exhibits one of the largest average differences in oxygen availability between tumor and adjacent healthy tissue^19^. We further define mechanistic insights governing this effect, showing that blockade of C1APP induction occurs independently of hypoxia-inducible factor (HIF)-1α or HIF-2α signaling and instead arises from arrest of C1APP protein translation. This translational arrest occurs downstream of the integrated stress response and is associated with impaired ribosome loading and sequestration of C1APP transcripts in hypoxia-associated stress granules. Importantly, we show that this hypoxia-induced defect is reversible by treatment with the RNA/DNA-integrating epitranscriptomic compound 5-azacytidine, but not by the closely related DNA-integrating epigenetic agent decitabine. Lastly, to see if C1APP components are excluded from hypoxic regions in human tumors, we analyzed expression patterns of IP subunits in nine stage II or stage III NSCLC patients and found significant regional exclusion from areas expressing the hypoxia marker carbonic anhydrase 9 (CA9) and found overall negative correlations between IP subunits and CA9 in six of seven patients with quantifiable CA9. Together, these findings establish that tumor hypoxia reversibly impairs C1APP induction and activity, defining a state of immunogenic dormancy and revealing a previously unrecognized role for stress granules in mediating immune escape in cancer.

### Hypoxia prevents augmented immunopeptide processing and presentation following IFN-γ

To ascertain the effect of hypoxia on cancer cell immunogenicity, we preconditioned human lung adenocarcinoma A549 cells to normal O_2_ (20% O_2_) or 2% O_2_ culture conditions for 48 hours before treatment with IFN-γ. HLA-presented immunopeptides were immunoprecipitated from cell lysates using the pan–HLA class I antibody W6/32, eluted, and analyzed by nanoLC–MS/MS using parallel accumulation–serial fragmentation (PASEF) with trapped ion mobility spectrometry and data-dependent acquisition (DDA-PASEF) on a Bruker timsTOF Pro 2. Eluted peptides were identified and quantified from the resulting spectra using a label-free immunopeptidomics workflow in Fragpipe^20,21^(See Methods, Extended Figure 1). Control groups included A549 cells preconditioned to normal O_2_ (∼20% O_2_; atmospheric oxygen) and/or cultures without IFN-γ treatment for a total of 4 experimental groups (n=3). Consistent with class I presentation, identified immunopeptides favored lengths of 9-10 amino acids, with a minority comprising longer lengths. Under normal O_2_, IFN-γ treatment significantly shifted HLA-immunopeptides to shorter lengths of 8 (p < 0.001) and 9 (p < 0.0001) amino acids. This was at the expense of longer amino acid lengths of 11-25 (all significantly reduced; p < 0.05 or lower). This shift was absent under hypoxia with the exception of a small increase in frequency of 8 amino acid-long immunopeptides (p < 0.05; Figure 1a), suggesting that antigen processing under normal O_2_ following IFN-γ treatment favors the creation of immunopeptides with optimal MHC class I length. Treatment with IFN-γ induced a significant increase in the number of unique immunopeptides (p < 0.01; Figure 1b), and the number of immunopeptides generated from common starting proteins (p < 0.0001; Figure 1c) under normal O_2_. Under hypoxia, fewer unique immunopeptides were identified at baseline (no IFN-γ treatment; p < 0.05), and fewer average immunopeptides per starting protein were observed (no IFN-γ treatment; p < 0.05), with no increases in either metric observed following IFN-γ treatment (Figure 1b-1c). In total, 18,936 immunopeptides were identified. Of these, 2,962 were common to all 4 experimental groups. Beyond these, less overlap was observed between experimental groups, with IFN-γ-treated cells under normal O_2_ displaying the largest set of unique immunopeptides (9,220). In comparison, only 917 unique immunopeptides were identified under hypoxia, irrespective of IFN-γ treatment (Figure 1d). To better visualize the effect IFN-γ has on immunopeptide presentation, we subtracted the normalized peak intensities of each IFN-γ-treated group from the corresponding untreated group and then sorted the peptides into 10 groups (Figure 1e). Notably, groups 1-3 represent over 73% of all immunopeptides, and are those whose frequencies benefitted from IFN-γ treatment under normal O_2_, but that were absent, reduced, or augmented less under hypoxia. In contrast, groups where peptides were increased by IFN-γ under hypoxia, but were absent, reduced, or increased less under normal O_2_ (groups 8-10), collectively represent less than 13% of identified immunopeptides. Principal component analyses reiterate these categorical findings, with hypoxic samples (irrespective of IFN-γ treatment) clustering together near normal O_2_ without IFN-γ, and only normal O_2_ with IFN-γ treatment displaying separation (Figure 1f). Together, these data indicate a clear benefit in immunopeptide processing and presentation following treatment with IFN-γ that is absent under hypoxia.

**Figure 1:**
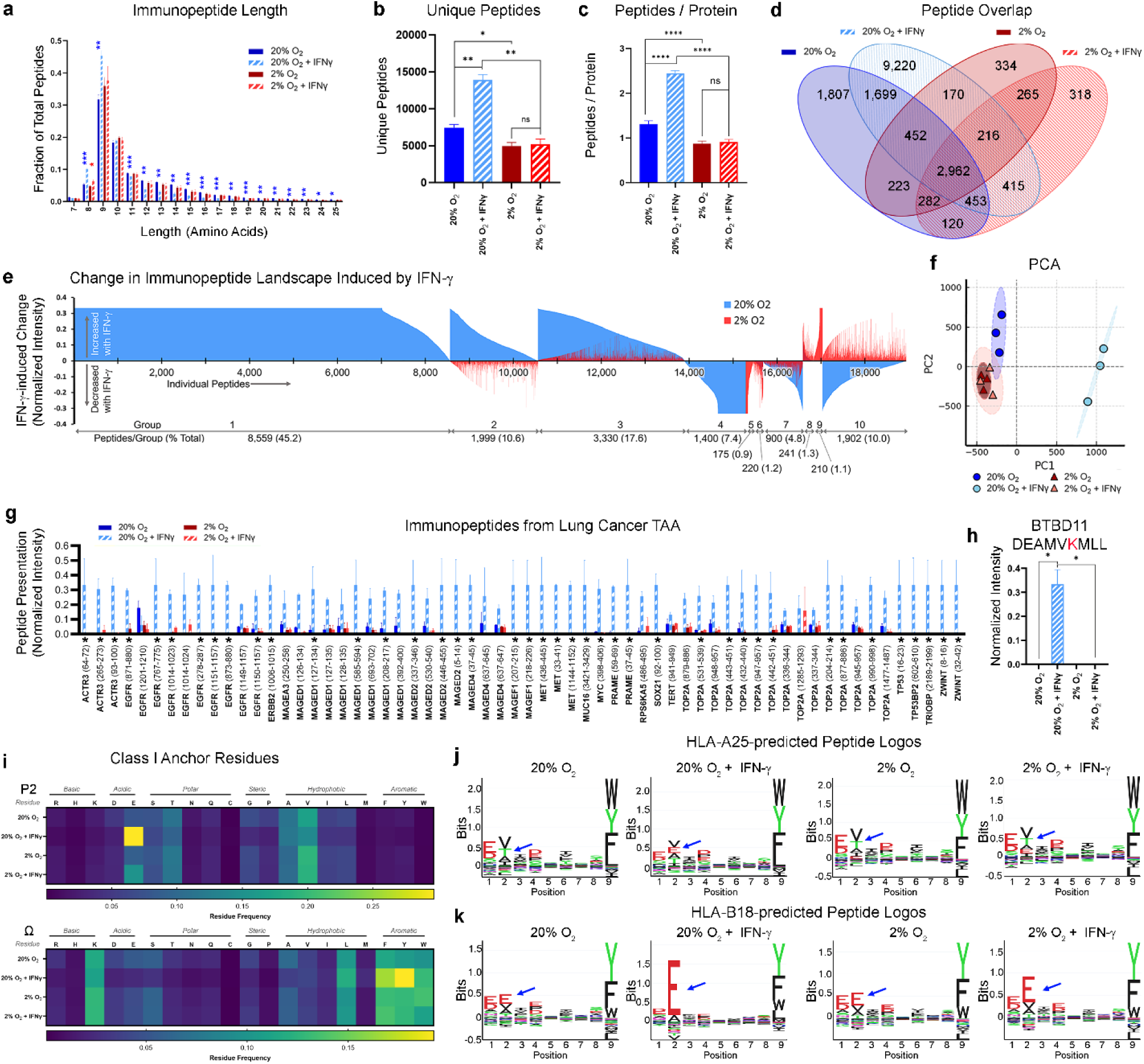
Hypoxia prevents augmentation of immunopeptide presentation following IFN-γ. **a,** Immunopeptidomic analysis was performed on HLA-immunoprecipitated eluates from A549 cells preconditioned in 20% O₂ or 2% O₂ for 48 h, followed by treatment with or without 10 ng/mL IFN-γ for an additional 24 h. Frequency distribution of peptide lengths for IFN-γ-treated versus untreated cells preconditioned in either 20% O₂ (blue asterisks) or 2% O₂ (red asterisks), assessed by Student’s t-test. **b**, The number of unique peptides identified per group. **c**, The average number of peptides derived from each source protein. **d**, The overlap of unique peptides between conditions. **e**, Landscape plot of all 18,902 identified peptides, displayed as the normalized intensity change induced by IFN-γ (treated – untreated) under 20% O₂ (blue) or 2% O₂ (red). Peptides were partitioned along the x-axis into 10 predefined categories: Group 1: observed only in 20% O₂ and increased by IFN-γ; Group 2: increased by IFN-γ in 20% O₂ but decreased in 2% O₂; Group 3: increased to a greater extent by IFN-γ in 20% O₂; Group 4: observed only in 20% O₂ without IFN-γ; Group 5: observed only in 2% O₂ without IFN-γ; Group 6: decreased to a greater extent by IFN-γ in 2% O₂; Group 7: decreased to a greater extent by IFN-γ in 20% O₂; Group 8: increased to a greater extent by IFN-γ in 2% O₂; Group 9: observed only in 2% O₂ + IFN-γ; Group 10: increased by IFN-γ in 2% O₂ but decreased in 20% O₂. Notably, Groups 1–3 represent peptides whose HLA presentation is enhanced by IFN-γ and collectively comprise >73% of all identified peptides. **f,** PCA of all identified peptides across conditions. **g,** Normalized intensities of immunopeptides derived from known lung cancer tumor-associated antigens. **h**, Normalized intensity of the BTBD11-derived neoantigen bearing a Q621K substitution (DEAMVKMLL) (mutation shown in red), quantified from inferred ion peaks. **i**, Amino acid frequencies at canonical HLA anchor positions (P2, top; PΩ, bottom) among all identified immunopeptides; Student’s t-test compares IFN-γ-treated versus untreated samples under 20% O₂ (blue asterisks) or 2% O₂ (red asterisk). **j, k,** Predicted peptide sequence logos for HLA-A25 (**j**) and HLA-B18 (**k**). Blue arrows highlight prominent amino acid changes corresponding to those in panel (**i**). For panels (**a–c**): *P≤0.05; **P≤0.01; ***P≤0.001; ****P≤0.0001 (Student’s t-test). For panel (**g**): *P≤0.05 for 20% O₂ vs 20% O₂ + IFN-γ <u>and</u> *P≤0.05 for 20% O₂ + IFN-γ vs 2% O₂ + IFN-γ. For panel (**i**): *P≤0.0001. n = 3 biological replicates for all panels (**a–k**). For **b-c**, bars represent mean ± s.e.m.

To focus on immunopeptides that might elicit an anti-tumor immune response, we plotted normalized peak intensities of immunopeptides mapping to known lung cancer TAA^22–29^, identifying 62 TAA-derived immunopeptides (Figure 1g). Of these, 41 (66%) displayed significantly increased intensities after IFN-γ treatment under normal O_2_, <u>and</u> significantly less intensities after IFN-γ treatment under hypoxia (p < 0.05 for both 20% O_2_ vs. 20% O_2_ + IFN-γ, <u>and</u> 20% O_2_ + IFN-γ vs. 2% O_2_ + IFN-γ). We were able to identify one neoantigen with repeatable detection, a BTBD11-derived neoantigen bearing a Q621K substitution (DEAMV**K**MLL) (Figure 1h). This neoantigen was only detected in IFN-γ-treated cells under normal O_2_ (Figure 1h; p <0.05 for 20% O_2_ vs. 20% O_2_ + IFN-γ and 20% O_2_ + IFN-γ vs. 2% O_2_ + IFN-γ).

To understand how hypoxia may affect IFN-γ-augmented immunopeptide composition, we quantified the frequency of amino acid frequencies in the second position (P2) and last position (Ω) (the quintessential anchor residue positions for HLA^30^), among unique peptides. P2 anchor residue composition differed significantly across experimental conditions (PERMANOVA on CLR-transformed frequencies, p = 0.017), driven largely by an increase in glutamate (E) frequency observed after IFN-γ treatment under normal O_2_, that is markedly attenuated under hypoxia. Ω anchor residue composition also differed significantly across experimental conditions (PERMANOVA on CLR-transformed frequencies, p = 0.011), driven by increases in the hydrophobic moiety leucine (L) and aromatic moieties phenylalanine (F), tyrosine (Y), and tryptophan (W) under normal O_2_, which were markedly attenuated under hypoxia (Figure 1i). Of these residues, changes in Ω-position L, F, Y, and W were identifiable in both unsupervised and allele-specific HLA-A and HLA-B peptide logo analyses (Figure 1j-1k, Extended Figures 2 and 3), whereas P2-position E was relegated to HLA-B alleles (Figure 1k, Extended Figures 2 and 3). Neither increase was observed in predicted immunopeptide logos following IFN-γ treatment under hypoxia (Figure 1j-1k). Interestingly, no discernable changes were observed in predicted immunopeptide logos between experimental groups for HLA-C alleles (Extended Figure 2). Collectively, these data indicate that the benefit IFN-γ bestows upon the number and variety of immunopeptides processed and presented by HLA are lost under hypoxic conditions, a phenomenon capable of affecting not only “normal self” immunopeptides, but those derived from TA and neoantigens as well.

### Hypoxia blocks cytokine-induced expression of C1APP proteins but not C1APP mRNA

Because we observed deficits in the landscape of immunopeptides presented by A549 cells under hypoxia, we examined C1APP protein expression (Figure 2a) in A549 lung adenocarcinoma preconditioned to normal O_2_ or hypoxia before IFN-γ treatment. Under hypoxia, HIF-1α and HIF-2α were stabilized, and hypoxia signaling components were largely unaffected by IFN-γ while conventional proteasome catalytic subunits and the alpha 3 (α3) subunit remained stable across oxygen conditions and IFN-γ doses. IFN-γ induced robust IP catalytic subunit β1i, β2i, and β5i expression under normal O_2_, whereas hypoxia markedly reduced or prevented induction of all three subunits (Figure 2b). Downstream of the IP, IFN-γ-induced increases in TAP1, tapasin, and calreticulin observed under normal O_2_ were strongly diminished under hypoxia. ERAP1 expression was largely maintained, while ERp57 showed a modest reduction (Figure 2c). Consistent with reduced C1APP induction, IFN-γ-driven upregulation of surface HLA was also significantly diminished under hypoxia, as assessed by flow cytometry (Extended Figure 4; Figure 2d). To ascertain if this phenomenon was limited to A549 cells, we performed similar experiments in 8 other epithelial-derived cancer cell lines, spanning human cancers: prostate, renal cell, hepatocellular, pancreatic, colorectal, melanoma, and finally in healthy (non-transformed) primary human lung epithelial cells, observing similar results (Extended Figure 5).

**Figure 2:**
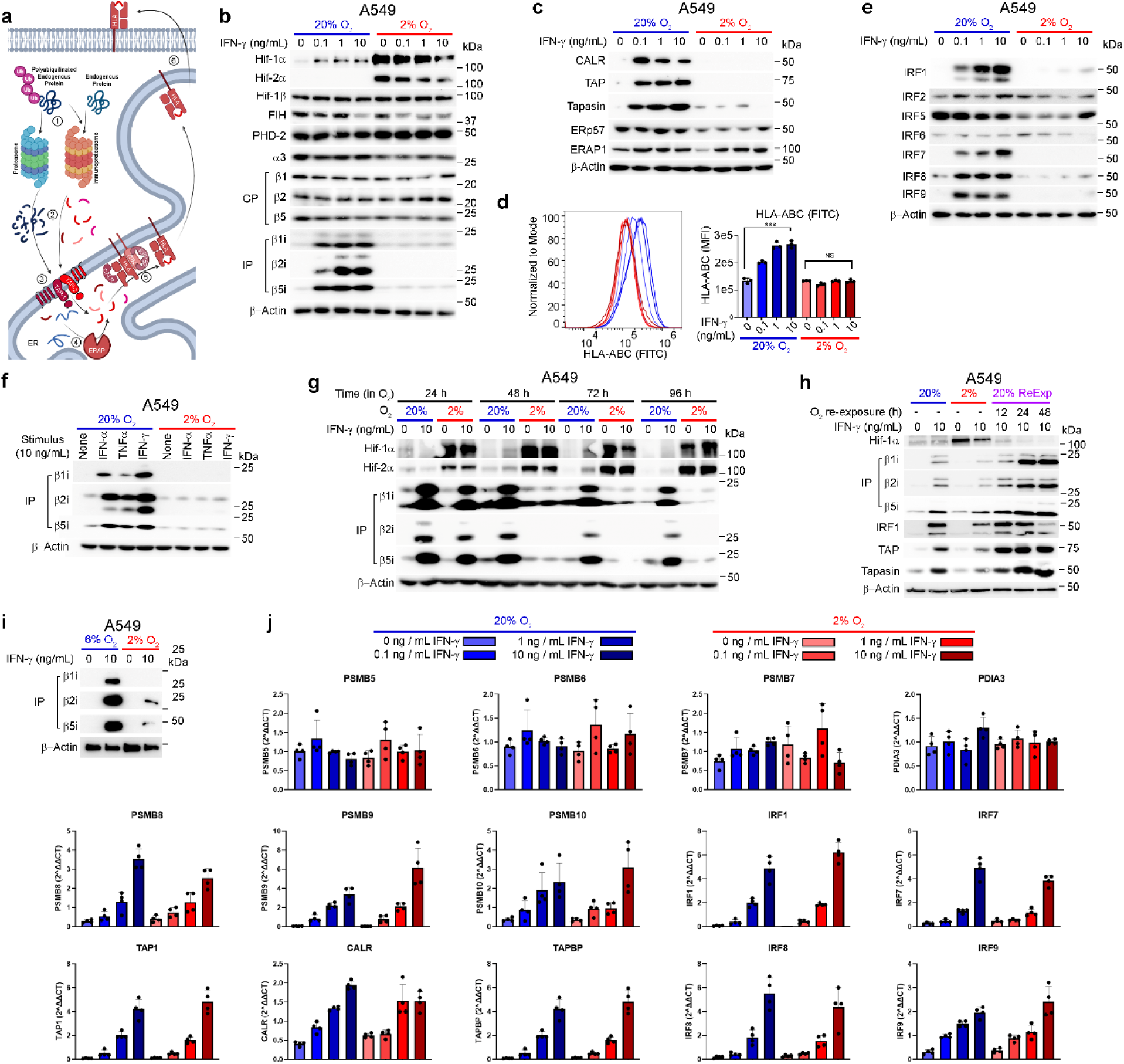
Hypoxia prevents the protein expression of the MHC class I antigen processing pathway. **a,** Schematic of the C1APP: ① Inflammatory cytokines induce the IP, which processes both ubiquitinated and non-ubiquitinated proteins and ② creates peptides with HLA-favoring C-termini, increasing HLA affinity and presentation likelihood. ③ Peptides are then transported into the ER via TAP (TAP-1/TAP-2), where ④ larger fragments may be further cleaved by ERAP. ⑤ ERp57 and calreticulin (CALR) associate with nascent MHC class I heavy chain–β2-microglobulin heterodimers together with tapasin to form the peptide-loading complex (PLC), which promotes the loading and stabilization of high-stability peptide–MHC class I complexes prior to ⑥ trafficking to the cell surface. Pathway components induced or up-regulated by IFN-γ are depicted in red tones. Graphics made in part using BioRender®. **b-c,** Immunoblot analysis of lysates from A549 cells preconditioned for 48 h to 20% O_2_ or 2% O_2_ before treatment with the indicated concentrations of IFN-γ for another 24 h, for the indicated HIF pathway proteins, conventional proteosome (CP) catalytic subunits, IP catalytic subunits (**b**), or extended MHC class I antigen processing pathway (C1APP) (**c**). **d**, Flow cytometry analysis of cell-surface HLA (pan HLA-ABC). Shown is a set of representative histograms (left) and aggregate MFI (right; n = 3). ***P≤0.001 (Student’s t-test). **e**, Immunoblot analysis of lysates from A549 cells as described in part **b** for IRF family member proteins. **f,** Immunoblot analysis of lysates from A549 cells preconditioned for 48 h to 20% O_2_ or 2% O_2_ before treatment with or without 10 ng/mL of the indicated cytokines for another 24 h for IP catalytic subunits. **g**, Immunoblot analysis of lysates from A549 cells preconditioned for the indicated times to 20% O_2_ or 2% O_2_ before treatment with or without 10 ng/mL of the IFN-γ for another 24 h for HIF-1α, HIF-2α, or IP catalytic subunits. **h**, Immunoblot analysis of lysates from A549 cells preconditioned for 48 h to 20% O2 or 2% O2 before treatment with or without 10 ng/mL of IFN-γ for another 24 h, or from A549 cells preconditioned for 48 h to 2% O2, and then re-exposed to 20% O2 for the indicated times along with 10 ng/mL IFN-γ. **i**, Immunoblot analysis of lysates from A549 cells preconditioned for 48 h to 20% O_2_ or 2% O_2_ before treatment with or without 10 ng/mL of the indicated cytokines. β-αctin is included as a loading control for **b-c** and **e-i**. **j**, RT-qPCR analysis of mRNA isolated from A549 cells (n=4) preconditioned for 48 h to 20% O_2_ or 2% O_2_ before treatment with the indicated concentrations of IFN-γ for another 24 h. Student’s t-tests were run between 20% and 2% O_2_ at each dose of IFN-γ; no significant differences were observed (n = 4). mRNA levels of conventional proteasome levels (PSMB5, PSMB6, PSMB7; not responsive to IFN-γ) are shown for context. Data presented in **b-i** are representative of at least three independent experiments. For **d** and **j**, bars represent mean ± s.e.m.

Because IFN-γ-responsive programs are coordinated by interferon regulatory factors (IRFs)^31^, we examined IRF protein expression. Induction of IRF1, IRF7, IRF8, and IRF9 was blocked under hypoxia, whereas IRF2, IRF5, and IRF6 were minimally effected (Figure 2e). IRF3 and IRF4 were not detected in A549 cells. To determine whether this effect was specific to IFN-γ, we stimulated cells with IFN-α or TNF-α, inflammatory cytokines also known to induce C1APP components^32,33^. Although both cytokines increased IP expression under normal O_2_, their ability to do so was similarly blocked under hypoxia (Figure 2f), suggesting this phenomenon is independent of stimulus type.

Time-course analyses showed that sustained hypoxia (≥48 h) was required to observe pronounced loss of inducible IP subunit expression (Figure 2g), whereas re-exposure to normal O_2_ after 48-hour hypoxic pre-conditioning, restored IFN-γ-induced IP induction within 12–24 h (Figure 2h). As a control validation, we repeated this experiment, comparing hypoxia-conditioned cells to cells pre-conditioned to 6% O₂, a physiologically relevant lung oxygen tension, and observed similar results (Figure 2i). In contrast to protein expression, RT–qPCR revealed comparable IFN-γ-induced expression of immunoproteasome and C1APP mRNA transcripts under normal O_2_ and hypoxia. We did not observe significant reductions in any of these mRNAs under hypoxia at any IFN-γ dose (Figure 2j), demonstrating that hypoxia limits inducible C1APP protein expression without reducing transcript abundance.

### Hypoxia-induced C1APP blockade is independent of IFN-γ signaling, autophagy, and HIF activity

To define the mechanism underlying hypoxia-induced C1APP blockade, we first assessed whether upstream IFN-γ signaling was impaired under hypoxia. IFN-γ stimulation induced robust phosphorylation of STAT1 under both normal O_2_ and hypoxia, indicating intact canonical signaling (Figure 3a). Consistent with this, surface expression of the IFN-γ receptor (IFNGR) was unchanged across oxygen conditions (Figure 3b). These findings indicate that proximal IFN-γ signaling remains intact under hypoxia.

**Figure 3:**
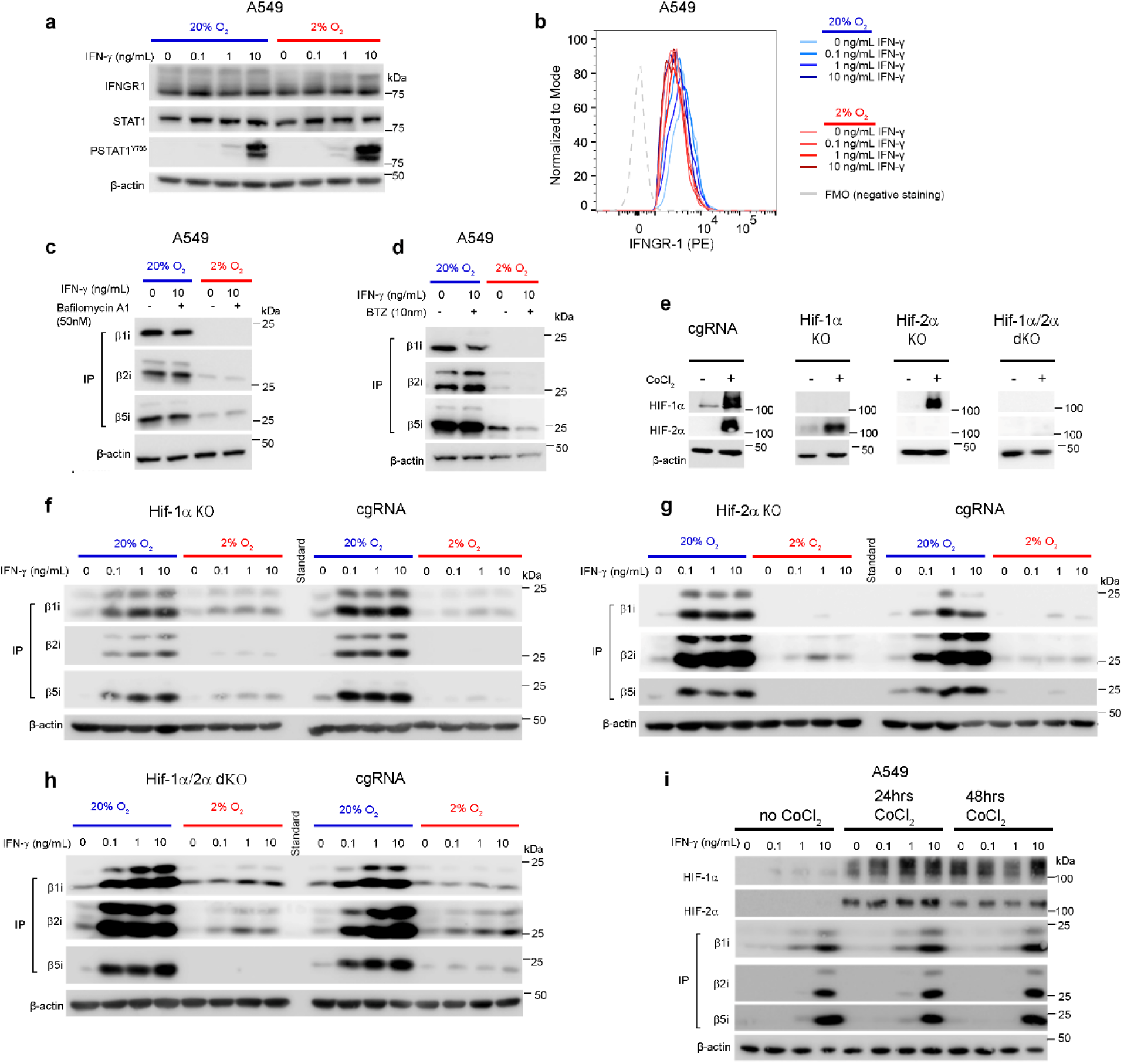
Hypoxia-induced blockade of C1APP proteins is HIF-independent and not attributed to protein degradation, or autophagy. **a**, Immunoblot analysis of lysates from A549 cells preconditioned for 48 h to 20% O_2_ or 2% O_2_ before treatment with the indicated concentrations of IFN-γ for another 24 h, for the indicated IFN-γ signaling components. **b**, Cell surface staining of IFNR-1, as measured by flow cytometry on A549 cells preconditioned for 48 h to 20% O_2_ or 2% O_2_ before treatment with the indicated concentrations of IFN-γ for another 24 h **c-d**, Immunoblot analysis of lysates from A549 cells preconditioned for 48 h to 20% O_2_ or 2% O_2_ along with or without 50 nM bafilomycin A1 (**c**) or 100 nM bortezomib (BTZ; **d**) for the indicated IP catalytic subunits. **e**, CRISPR/Cas9 was used to generate HIF-1α, HIF-2α, or HIF-1α/HIF-2α double knockout A549 clones. CoCl_2_ treatment was then used to chemically stabilize HIFs, and immunoblotting was performed to assess HIF KO status**. f-h**, A549 HIF KO cells were then used in experimental setups as described in panel **(a)**, along with control guide RNA (cgRNA) CRISPR/Cas9 controls. Immunoblotting was used to assess expression levels of the indicated IP catalytic subunits using HIF-1α KO (**f**), HIF-2α KO (**g**) or HIF-1α/HIF-2α double KO A549 cells (**h**). **i**, CoCl_2_ treatment was used to chemically stabilize HIFs for 24 or 48 hours before treatment with the indicated concentrations of IFN-γ for another 24 h Immunoblotting was used to assess expression levels of the indicated IP catalytic subunits.

Previous studies have implicated severe hypoxia-associated autophagy in the regulation of baseline C1APP expression in colorectal cancer cells^17^. To test this possibility, we treated hypoxic A549 cells with the lysosomal inhibitor bafilomycin A1 to block autophagy-mediated degradation. Inhibition of autophagy failed to restore IFN-γ-induced immunoproteasome expression under hypoxia (Figure 3c), arguing against autophagy as the primary cause of reduced C1APP protein levels at these oxygen levels.

Because transcription of immunoproteasome and C1APP genes was not lost under hypoxia, we asked whether ubiquitin-proteasome system (UPS) degradation was preventing stable expression of these transcripts via accelerated protein turnover. To test this, we treated normal O_2_ or hypoxia preconditioned A549 cells with the proteasome inhibitor bortezomib (BTZ) to inhibit UPS-directed protein turnover. Inhibition of proteasome function failed to enhance IFN-γ-induced IP subunit expression under hypoxia, suggesting that similar to autophagy, reduced C1APP protein induction is independent of UPS degradation under hypoxia (Figure 3d).

We next investigated the role of hypoxia-inducible factors (HIFs), which mediate canonical transcriptional responses to reduced oxygen availability^34^. CRISPR/Cas9–mediated knockout (KO) of HIF-1α, HIF-2α, or combined HIF-1α/HIF-2α (Figure 3e) did not restore IFN-γ-induced immunoproteasome expression under hypoxia (Figure 3f-3h). To further assess whether HIF activation is sufficient to suppress inducible C1APP expression, we treated A549 cells under normal O_2_ with cobalt chloride, a pharmacologic hypoxia mimetic that stabilizes HIF proteins independently of oxygen tension. Cobalt chloride treatment did not phenocopy hypoxia-induced C1APP suppression (Figure 3i). Together, these data indicate that hypoxia-induced reduction of inducible C1APP protein expression occurs independently of IFN-γ signaling, autophagy-or UPS-mediated protein degradation, and canonical HIF-1α/2α–dependent pathways.

### 5-azacytidine prevents hypoxia-induced C1APP blockade and restores IFN-γ-augmented unique immunopeptide numbers and diversity

Given that hypoxia-induced loss of C1APP protein expression occurs despite intact transcriptional responses to IFN-γ, we next asked whether this phenotype could be modulated pharmacologically. 5-azacytidine (5-AZA) and decitabine (5-aza-2′-deoxycytidine; DAC) are closely related, first-generation epigenetic therapeutics that differ primarily in their capacity for RNA versus DNA incorporation. 5-AZA is a ribonucleoside analog that incorporates predominantly into RNA (approximately 80-90%) with a smaller fraction incorporating into DNA (approximately 10-20%), whereas DAC, lacking a ribose 2′-hydroxyl group, incorporates exclusively into DNA^35^. Although both agents induce DNA hypomethylation through trapping of DNA methyltransferases, only 5-AZA substantially perturbs RNA methylation by engaging cytosine-5 RNA methyltransferases. Since both 5-AZA and DAC have reported ability to up-regulate the C1APP in different contexts^36,37^, we assessed whether their administration could prevent hypoxia-induced C1APP blockade. Treatment with 5-AZA during hypoxic pre-conditioning preserved IFN-γ-induced expression of immunoproteasome catalytic subunits (Figure 4a). In contrast, treatment with the structurally related analog decitabine (5-aza-2′-deoxycytidine; DAC), which incorporates exclusively into DNA, failed to prevent hypoxia-associated loss of C1APP protein expression (Figure 4b). This distinction is consistent with RNA-associated, rather than DNA-restricted, effects in preserving inducible C1APP protein expression under hypoxia.

**Figure 4:**
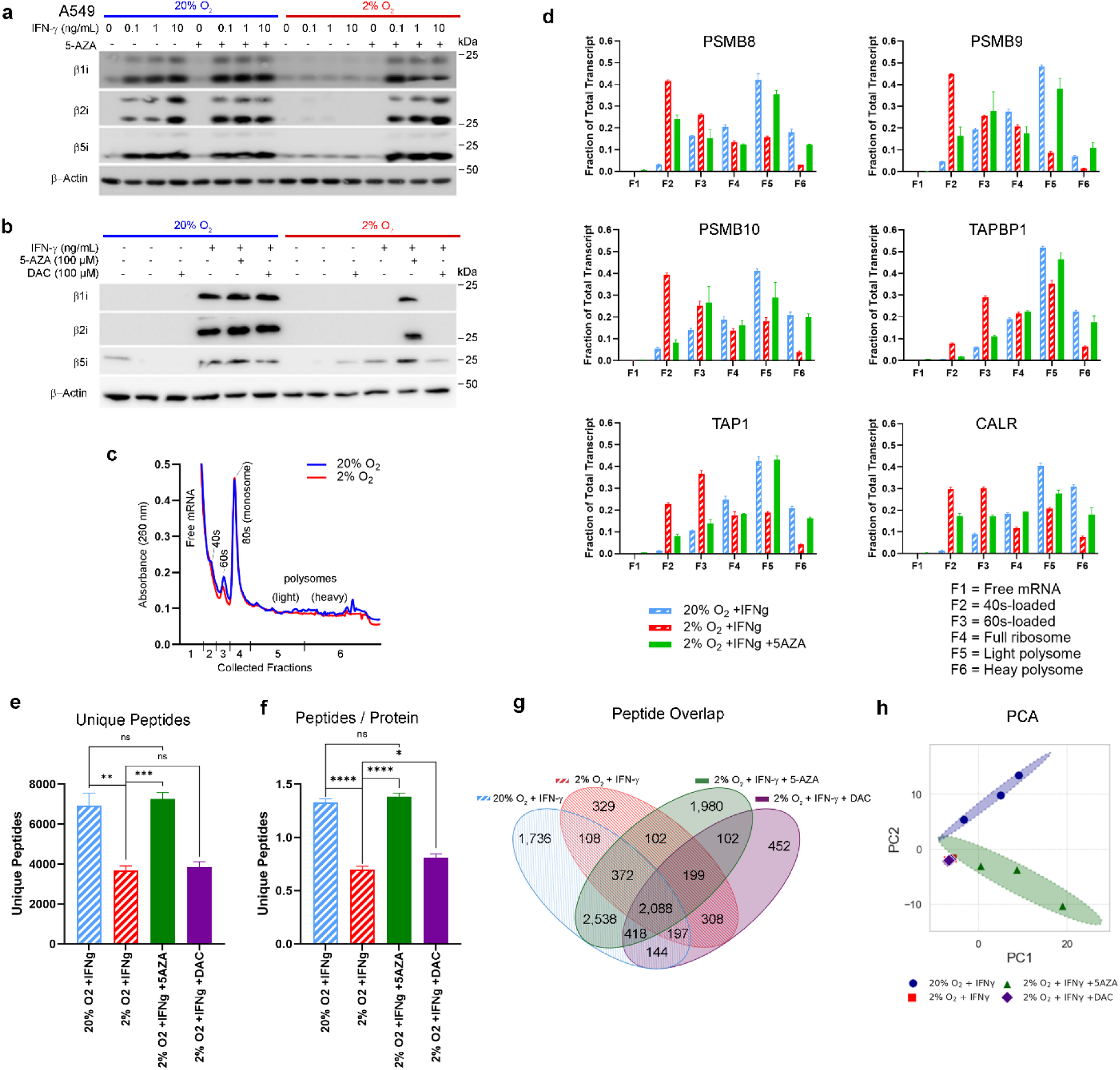
5-azacytidine (5-AZA), but not decitabine (DAC), restores immunopeptide presentation following IFN-γ, and prevents protein translation arrest of the C1APP mRNAs. **a**, Immunoblot analysis of lysates from A549 cells preconditioned to 20% O_2_ or 2% O_2_ for 48 h with or without 5-AZA followed by treatment with the indicated dose of IFN-γ for an additional 24 h for the indicated IP catalytic subunits. **b**, Immunoblot analysis of lysates from A549 cells preconditioned to 20% O_2_ or 2% O_2_ for 48 h with or without 5-AZA or DAC before overnight stimulation with 10 ng/mL IFN-γ for the indicated IP catalytic subunits. β-actin is included as a loading control for **a** and **b. c, d**, Polysome profiling was performed on A549 cells preconditioned to 20% O_2_ or 2% O_2_ for 48 h with or without 5-AZA followed by treatment with or without 10 ng/mL IFN-γ for an additional 24 h. Representative polysome traces are shown (**c**). Fractions were pooled, approximating free mRNA (F1), 40s-bound mRNAs (F2), 60s-bound mRNAs (F3), monosome-bound mRNAs (F4), light polysome-bound mRNAs (F5), and heavy polysome-bound mRNAs (F6). RT-qPCR was then performed for the indicated IP catalytic subunits or the C1APP members demonstrating protein blockade in Figure 2 (**d**). Transcript levels are displayed as the relative transcript abundance within each pooled polysome fraction (F1-F6). Error bars are +/-SEM from three technical replicates. Shown is a representative experiment of 4 repeats. Note the shift to pre-monosome fractions under hypoxic culture conditions, and normalization of polysome fractions F5 and F6 with 5-AZA. **e-h**, Immunopeptidomic analysis was performed on HLA-immunoprecipitated eluates from A549 cells preconditioned in 20% O₂ or 2% O₂ along with or without 5-AZA or DAC for 48 h, followed by treatment with or without 10 ng/mL IFN-γ for an additional 24 h. Shown are: the number of unique peptides identified (**e**), the average number of peptides derived from each source protein (**f**), the overlap of unique peptides between conditions (**g**), and PCA of all identified peptides across conditions (**h**). n = 3 and **P≤0.01; ***P≤0.001 for **e-h**. For **d**, **e**, and **f**, bars represent mean ± s.e.m.

Because C1APP transcripts remain inducible under hypoxia despite blocked protein expression, we next examined whether hypoxia alters translation of these transcripts using polysome profiling. Overall, polysome traces were similar between O_2_ conditions, with negligible reductions in heavy and light polysome fractions under hypoxia (Figure 4c). We then pooled fractions approximating free mRNA (F1), 40s-loaded mRNA (F2), 60s-loaded mRNA (F3), monosome-loaded mRNA (F4), light polysome-loaded mRNA (F5), and heavy polysome-loaded mRNA (F6), and conducted RT-qPCR for mRNAs corresponding to C1APP proteins blocked under hypoxia. Under normal O_2_, the majority of C1APP mRNAs were found in polysome fractions (blue bars; F5-F6). Critically, a shift toward pre-monosome fractions (red bars; F2 and F3) were observed for C1APP mRNAs under hypoxia, suggesting protein translation arrest for these mRNAs. Inclusion of 5-AZA (green bars) in hypoxic preconditioning normalized C1APP protein translation (Figure 4d), consistent with observed protein induction (Figure 4a-4b). Overall, these findings indicate that hypoxia limits inducible C1APP expression at the level of mRNA utilization rather than transcription, and that this phenomenon can be prevented by 5-AZA.

To determine whether preservation of C1APP protein expression by 5-AZA translated into functional antigen presentation, we assessed HLA-presented immunopeptides under hypoxia along with 5-AZA or DAC (Extended Figure 6). As before, hypoxia significantly reduced the number of unique immunopeptides, and the diversity of immunopeptides presented by HLA (p < 0.01 for both). Inclusion of 5-AZA significantly increased the number of unique immunopeptides and diversity of immunopeptides relative to untreated hypoxic controls (p < 0.001 and p < 0.01 respectively), restoring levels comparable to those observed following IFN-γ treatment under normal O_2_ (Figure 4e-4f), as well as IFN-γ-favored anchor residues like P2-position glutamine for HLA-B18 and HLA-B44 (Extended Figures 7 and 8). Many 5-AZA augmented peptides overlapped with those found under normal O_2_, but many were unique to 5-AZA-treated samples (Figure 4g), likely reflecting the large number of gene expression changes known to be induced by 5-AZA^36,38^. Principle component analysis reflects these overlaps, with IFN-γ and DAC-treated hypoxic samples overlapping with IFN-γ (only)-treated hypoxic samples, while normal O_2_ IFN-γ-treated samples and hypoxic IFN-γ and 5-AZA treated samples displaying separation (Figure 4h).

Because 5-AZA, but not DAC, prevented hypoxia-induced blockade C1APP proteins, we asked whether hypoxia alters m5C patterns on C1APP mRNAs in a manner consistent with a direct, transcript-centric mechanism. To test this, we performed nanopore direct RNA sequencing to generate predicted m5C modification calls, and m5C methylated RNA immunoprecipitation sequencing (MeRIP-seq) in A549 cells preconditioned to normal O_2_ or hypoxia for 48 hours prior to IFN-γ treatment. Across covered sites within C1APP transcripts, nanopore analyses did not identify reproducible hypoxia-associated differences in predicted m5C modification calls. Consistent with this, m5C MeRIP-seq did not reveal significant m5C peak enrichment or differential m5C peak signal in C1APP mRNAs under hypoxia. At the transcriptome level, differential peak analyses likewise did not yield significantly enriched pathways or ontologies after multiple-testing correction that would explain the C1APP translational arrest (Extended Figures 9 and 10). While these findings do not rule out a role for m5C in hypoxia-induced C1APP blockade, they indicate that direct hypomethylation of C1APP mRNA transcripts or stable transcriptome-wide changes in mRNA m5C may not account for the observed effects of 5-AZA at the tested time or hypoxic conditions.

### IFN-γ-induced immunoproteasome subunit mRNAs are sequestered in hypoxia-associated stress granules

Hypoxia is a known trigger of the integrated stress response (ISR), which can impair protein translation^39^ and sequester nascent mRNA in stress granules, both stabilizing them, but precluding their translation^40^. Multiple pathways engage the ISR, but all lead to phosphorylation of the translation initiation factor 2 α (eIF2α), reducing the activity of the eIF2 complex, tempering protein translation^39^. Since 5-AZA prevents hypoxia-induced C1APP protein translation arrest, we asked if 5-AZA can also prevent hypoxia-induced integrated stress response. Consistent with previous reports^39^, hypoxia induced the phosphorylation of eIF2α in IFN-γ-treated hypoxic A549 cells indicating engagement of the ISR. Inclusion of 4 μM 5-AZA (or greater) reduced or prevented ISR engagement and restored IFN-γ induction of immunoproteasome subunits (Figure 5a). To assess stress granule induction, A549 cells preconditioned to normal O_2_ or hypoxia for 48 hours before IFN-γ treatment, were immunofluorescently stained for G3BP1, a well-established stress granule marker^41^. Sodium arsonite (NaAsO_2_) is a potent inducer of the ISR and commonly used positive control for stress granule induction^41^. While readily detectable, G3BP1 staining in IFN-γ-treated A549 cells under normal O_2_ is diffuse and cytoplasmic. In contrast, treatment with NaAsO_2_, or preconditioning to hypoxia induced robust formation of stress granules as indicated by punctate G3BP1 staining. Consistent with C1APP protein translation and expression patterns (Figure 4a-4b; 4h), inclusion of 5-AZA, but not DAC, during hypoxia preconditioning prevented stress granule formation (Figure 5b). To deduce if these stress granules contain C1APP mRNAs, we performed RNA fluorescence *in situ* hybridization (RNA-FISH) to detect PSMB8 (β5i), PSMB9 (β1i), and PSMB10 (β2i) mRNAs, and found strong mRNA hybridization signals for each in hypoxia-associated stress granules (Figure 5c-5e). Lastly, we asked if NaAsO_2_ can phenocopy hypoxia-induced blockade of the C1APP in normal O_2_, observing greatly diminished C1APP induction following IFN-γ with NaAsO_2_ treatment (Figure 5f), suggesting induction of ISR/stress granules is sufficient to arrest C1APP protein translation.

**Figure 5:**
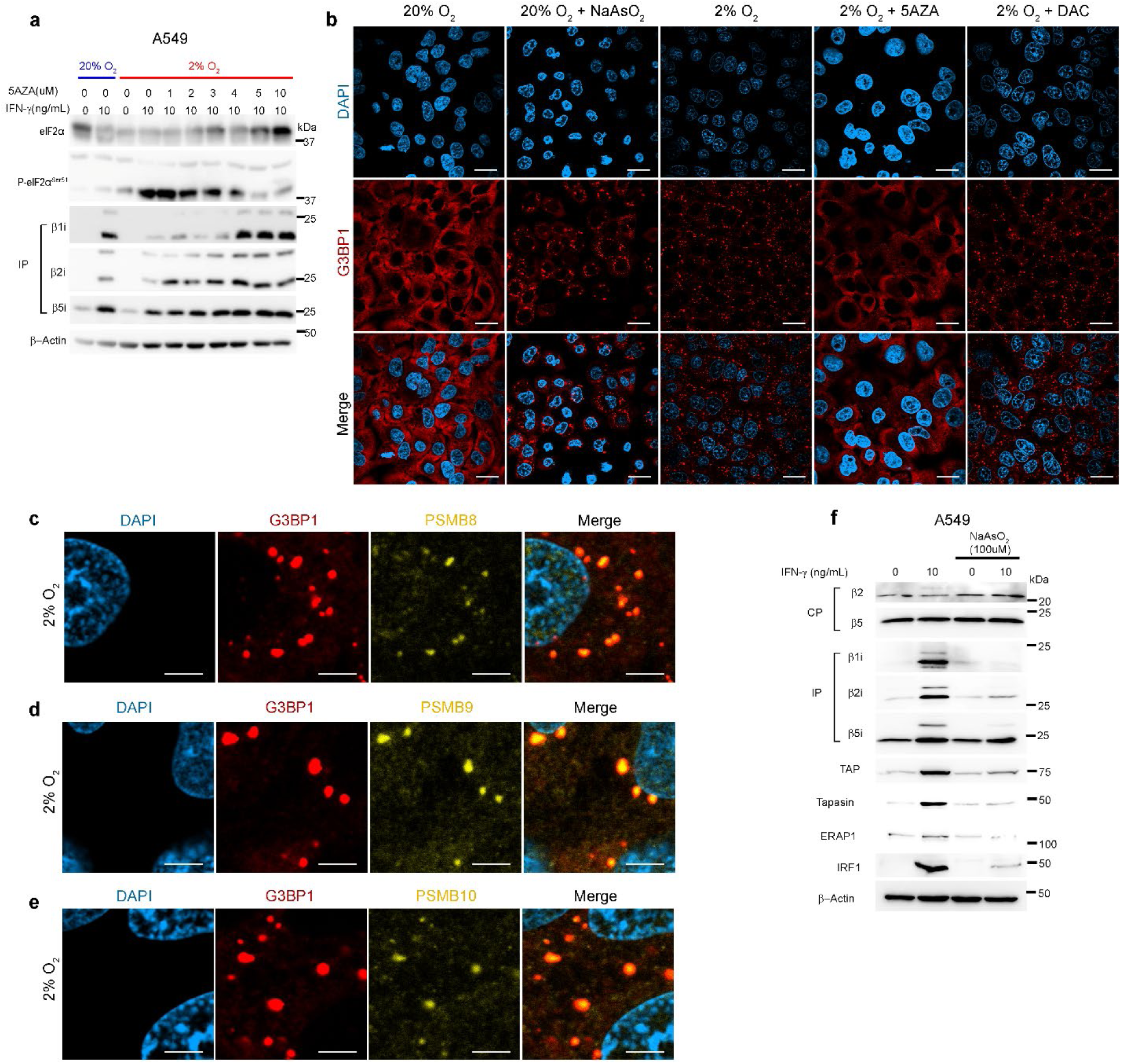
Hypoxia-induced stress granules sequester C1APP mRNA and prevent expression of C1APP proteins. **a**, Immunoblot analysis of lysates from A549 cells preconditioned to 20% O_2_ or 2% O_2_ for 48 h with or without the indicated dose of 5-AZA before 24 h stimulation with or without 24 h IFN-γ for total eIF2α, phospho-eIF2α, or indicated IP catalytic subunits. **b,** Representative images of immunofluorescence staining for stress granule marker G3BP1. A549 cells were preconditioned to 20% O_2_ or 2% O_2_ for 48 h with or without the indicated compounds before overnight stimulation with IFN-γ. Nuclei were counterstained with DAPI, and merged images were generated from all three channels. Scale bars = 20 μm. **c-e**, Representative images of fluorescent in situ hybridization (FISH) for PSMB8 (**c**), PSMB9 (**d**), or PSMB10 (**e**) along with immunofluorescent (IF) staining for G3BP1 in A549 cells preconditioned to 20% O_2_ or 2% O_2_ for 48 h with or without the indicated compounds before overnight stimulation with IFN-γ. Nuclei were counterstained with DAPI, and merged images were generated from all three channels. Scale bars = 5 μm. **f**, Immunoblot analysis of lysates from A549 cells treated with or without NaAsO_2_ during stimulation with or without IFN-γ for 24 h, for the indicated CP catalytic subunits, IP catalytic subunits, or extended C1APP proteins. NaAsO_2_ was added one hour following addition of IFN-γ. β-actin is included as a loading control for **a** and **f.**

### Immunoproteasome subunits are excluded from hypoxic regions in NSCLC tumors

Because C1APP protein translation is blocked under hypoxia, we next asked if C1APP expression is excluded from hypoxic tumor regions in human patients. To answer this, we performed immunofluorescent staining for the C1APP proteins β1i, β2i, or β5i, along with the hypoxia marker carbonic anhydrase 9 (CA9) in formalin-fixed paraffin embedded tissue sections containing grade II or grade III NSCLC tumors from nine patients (Figure 6a-6e). To quantify regional co-localization and regional exclusion between each immunoproteasome subunit and CA9, we computationally-derived superpixels in tiled images spanning large cross-sections of the tumor bed and analyzed threshold-defined quadrants of super pixels containing strong CA9 signal only (CA9^+^ alone), immunoproteasome only (β1i^+^, β2i^+^, or β5i^+^ alone), neither (double negative), or both (co-regional) (see Methods). For each immunoproteasome subunit, we found significantly less co-regional superpixels versus those with CA9^+^ alone (p < 0.05 for each) and versus those with β1i^+^ (p < 0.001), β2i^+^ (p < 0.001), or β5i^+^ alone (p < 0.0001) indicating large amounts of regional mutual exclusion (Figure 6b, 6d, 6f). As a complementary approach, we also calculated Spearman correlation coefficients between each immunoproteasome subunit and CA9 fluorescent signals in seven out of nine patients displaying at least 1% CA9^+^ tumor surface area and found significant negative correlations in two or all three immunoproteasome subunits in six patients (Figure 6g). Taken together, these data suggest the C1APP proteins β1i, β2i, or β5i are excluded from hypoxic regions in the majority of NSCLC patients and raise the likelihood that hypoxia-induced C1APP blockade diminishes immunogenicity in human tumors.

**Figure 6:**
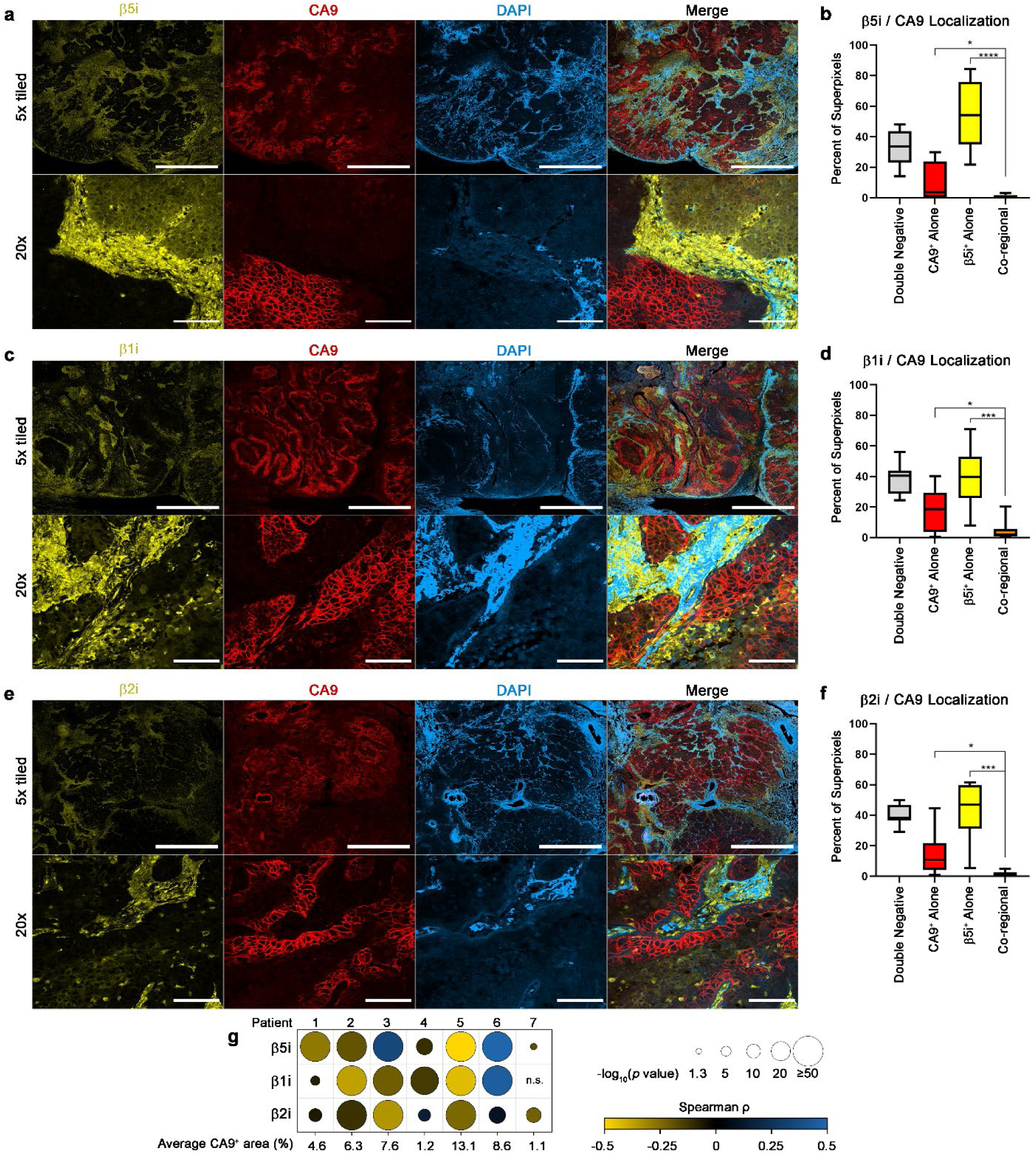
Catalytic immunoproteasome subunits are spatially excluded from hypoxic regions in human NSCLC tumors. **a-f**, Representative images of fluorescent staining for β5i (**a**), β1i (**c**), and β2i (**e**) along with immunofluorescent staining for CA9 of human NSCLC tumors (n=9). DAPI counterstain also includes bright autofluorescent signal from RBCs demarcating vasculature. Low powered (5x) tile images were stitched together (top; 3 x 4 tiles; scale bar = 2 mm), while a single representative higher powered (20x) image is shown (bottom; scale bar = 100 μm). Superpixels were computationally generated within defined tumor regions, and median fluorescence intensities of β5i (**b**), β1i (**d**), or β2i (**f**) and CA9 channels were quantified per superpixel to evaluate exclusive versus co-regional staining patterns. *P≤0.05; ***P≤0.001; ****P≤0.0001 (Student’s t-test; n = 9). **g**, Featured heatmap of Spearman correlation coefficients (with circle sizes representing p values) between CA9 and each IP subunit across tumor sample superpixels with at least 1% average CA9^+^ area (n = 7). n.s. = not significant. For **b**, **d**, and **f**, bars represent mean ± s.e.m.

## Discussion

Adaptive anti-tumor immunity ultimately depends on a non-negotiable requirement: tumor cells must process and present antigen on MHC class I molecules. Even when tumor-reactive CD8⁺ T cells are successfully primed and expanded, failure of tumor cells to display cognate antigen represents a terminal bottleneck that precludes immune recognition^10^. Tumor loss of the C1APP (including HLA) represents one of the longest-known modalities of tumor immune escape, for which few intrinsic mechanisms have been identified explaining its loss^42–45^. Here, we identify tumor hypoxia as a potent and reversible constraint on inducible C1APP, revealing a mechanism by which tumors can remain immunologically silent despite intact inflammatory signaling, effectively becoming “immunogenically dormant.”

A central implication of these findings is the breadth of antigenic loss imposed by hypoxia. Hypoxic conditions prevent IFN-γ-mediated expansion of the immunopeptidome, reducing both the number and diversity of HLA-presented peptides. We show this effect can extend to immunopeptides derived from tumor-associated antigens as well neoantigens, indicating that hypoxia broadly suppresses tumor immunogenicity rather than selectively reshaping epitope hierarchies. By insulating tumor cells from immune-derived cytokine cues, hypoxia disconnects inflammatory signaling from antigen presentation, thereby limiting immune recognition independently of antigen availability, mutational burden, or defects in T cell priming.

The relevance of this mechanism is amplified by the prevalence and heterogeneity of hypoxia within solid tumors. Oxygen deprivation is spatially and temporally dynamic, fluctuating with vascular instability and metabolic demand^46^. NSCLC, in particular, exhibits pronounced oxygen gradients both between tumor and adjacent normal tissue and within individual lesions^19^. Our observation that immunoproteasome expression is largely excluded from hypoxic tumor regions in advanced NSCLC specimens supports the idea that hypoxia imposes regional constraints on tumor immunogenicity in human patients. Such compartmentalization may help explain heterogeneous immune infiltration and incomplete therapeutic responses within otherwise immune-engaged tumors^47^.

These findings extend recent work demonstrating that near-anoxic stress suppresses baseline MHC class I abundance through hypoxia-dependent autophagy and lysosomal degradation^17^. In contrast to that paradigm of constitutive MHC-I turnover under extreme oxygen deprivation, our data identify a translational checkpoint that selectively impairs cytokine-driven induction of the C1APP at physiologically relevant hypoxic tensions. Because oxygen levels around the 2% range are common across solid tumors and span broad intratumoral territories^48^, this mechanism may operate across larger tumor compartments and across a wider fraction of patients than processes confined to near-anoxic niches. This distinction is particularly consequential in epithelial-derived cancers, where basal C1APP expression is low and immune visibility depends on IFN-γ-mediated upregulation^33^. The absence of autophagy involvement in our system, together with preserved cytokine signaling and transcriptional induction, underscores the existence of distinct, oxygen-dependent modes of antigen presentation failure that likely operate across different hypoxic niches. Because inducible C1APP governs the emergence of neoantigen- and TAA-derived epitopes under inflammatory pressure, its translational suppression may disproportionately impair adaptive anti-tumor immunity, even in regions that retain detectable baseline MHC-I expression.

Mechanistically, our data point to stress-responsive translational control as a key feature of hypoxia-induced C1APP blockade. Under hypoxia, C1APP transcripts remain inducible but fail to efficiently engage translating ribosomes and associate with hypoxia-induced stress granules. Whether stress granules actively enforce translational arrest or arise because of it remains unresolved^49^, and our data do not assign causality. Rather, they position C1APP transcripts within a hypoxia-responsive translational program that is incompatible with rapid protein induction, linking microenvironmental stress to impaired immune visibility.

The ability of 5-AZA to preserve inducible C1APP expression and restore immunopeptide diversity under hypoxia further supports a translationally regulated mechanism. Although we did not observe stable hypoxia-associated changes in m5C methylation across the transcriptome, these data do not exclude transient or context-dependent roles for RNA modifications. Notably, 5-AZA has well-described non-canonical effects on translation and stress responses^50^, emphasizing that its activity is dose- and context-dependent. Within this framework, our findings are most consistent with modulation of stress-adaptive translational programs rather than reversal of a fixed epitranscriptomic lesion.

Together, these data identify hypoxia-induced, stress-associated translational repression of the C1APP as an underappreciated mechanism of tumor immune escape and highlight stress-responsive translation as a critical layer of immune regulation within the tumor microenvironment.

## Supporting information

All extended data figures, legends, and supplementary data

## Methods

### Cell lines, cell culture, and treatments

A549, HEPG2, SKMEL2, A498, DU145, PANC1, HCT116, and 786-O are originally from the American Type Culture Collection (ATCC) authenticated by STR profiling (ATCC reference profiles). All cell lines were regularly verified as mycoplasma-free using PCR-based detection methods. Cell lines were cultured in low Glucose Dulbecco’s modified Eagle’s Medium (DMEM; Gibco) supplemented with 10% (v/v) heat inactivated fetal bovine serum (Gibco), penicillin (100 U/ml), streptomycin (100 μg/ml) (Gibco), 1% (v/v) 1 M HEPES (Gibco) and 1% (v/v) MEM non-essential amino acids (Corning). Cells were maintained at 37°C in a humidified incubator (HERAell VIOS) with 5% (v/v) CO_2_ and passaged at approximately 70-80% confluence. Hypoxic conditions were achieved by incubating cells in a dedicated humidified incubator with 5% CO_2_ (v/v) and 2% O_2_ (maintained by continuous N_2_ displacement). Primary human small airway epithelial cells (HSAEC; ATCC PCS-301-010) were cultured in T-25 tissue culture flasks using Airway Epithelial Cell Basal Medium (ATCC PCS-300-030) supplemented with the Bronchial Epithelial Cell Growth Kit (ATCC PCS-300-040), as recommended by the supplier. Primary human lung epithelial cells were similarly passaged at 70-80% confluence. Low passage numbers of primary lung epithelial cells (≤ passage 5) were used for experiments. Where indicated, cells were treated with 5-azacytidine (5-AZA), 5-aza-2′-deoxycytidine (DAC), bafilomycin A1, bortezomib, recombinant human IFN-γ, recombinant human IFN-α, recombinant human TNF-α, cobalt chloride (CoCl₂), or sodium arsenite (NaAsO₂) at the indicated concentrations and durations. For all treatments, reagents were freshly prepared as 1,000× working solutions and diluted in dPBS immediately prior to a final 1:1,000 dilution into culture medium, such that the vehicle volume constituted 0.1% of the total culture volume across all experimental conditions and vessel formats. Vehicle-only controls were included where appropriate.

### Western blot / Immunoblot

Whole-cell lysates were prepared on ice using radioimmunoprecipitation assay (RIPA) buffer (Thermo Fisher Scientific) supplemented immediately prior to use with protease and phosphatase inhibitor cocktails (Thermo Fisher Scientific). Cells were washed with ice-cold phosphate-buffered saline (PBS), lysed in RIPA buffer, and incubated on ice for 30 min with intermittent mixing. Lysates were clarified by centrifugation at 12,500 rpm for 20 min at 4 °C, and supernatants were collected for downstream analysis. Protein concentrations were determined using the DC Protein Assay (Bio-Rad), and samples were normalized to equal protein concentrations in RIPA buffer.

For SDS–PAGE, 10–30 µg of total protein per sample was denatured in 1× Laemmli sample buffer containing β-mercaptoethanol at 95 °C for 5 min and resolved on polyacrylamide gels under denaturing conditions. Proteins were transferred onto 0.2-µm polyvinylidene difluoride (PVDF) membranes (Bio-Rad) using a semi-dry transfer system operated under constant current conditions.

Membranes were blocked in commercial blocking buffer (Bio-Rad) for 30 min at room temperature and incubated with primary antibodies (Supplementary Table 1) diluted in blocking buffer overnight at 4 °C with gentle agitation. Membranes were washed five times with Tris-buffered saline containing 0.1% Tween-20 (TBST) and then incubated with species-appropriate horseradish peroxidase (HRP)–conjugated secondary antibodies for 1 h at room temperature. Following five additional washes in TBST, protein bands were visualized by enhanced chemiluminescence using Clarity Max Western ECL Substrate (Bio-Rad) and detected by autoradiography or digital imaging.

### Flow cytometry

Non-specific antibody binding was blocked by incubating cells for 30 min at 4 °C in PBS containing 2% fetal bovine serum (FBS), using 300 µL per 1.0 × 10⁶ cells. Cell-surface staining was performed by incubating cells for 30 min at 4 °C with FITC-conjugated anti–HLA-ABC or PE-conjugated anti-IFNGR-1antibodies (Supplementary Table 1). Cells were subsequently washed with PBS and resuspended in PBS containing 2% bovine serum albumin (BSA) and 7-aminoactinomycin D (7-AAD) for viability discrimination prior to acquisition.

Data were acquired on a BD LSR II flow cytometer equipped with a custom optical configuration at the Carver College of Medicine Flow Cytometry Core Facility. Control samples included unstained cells, single-color compensation controls, and fluorescence-minus-one (FMO) controls. Compensation matrices were calculated at the time of acquisition using BD FACSDiva software. Data were analyzed using FlowJo software v10.10 (Becton, Dickinson and Company) using a sequential, gate-based strategy as exemplified in Extended Figure 4.

### RNA isolation, reverse transcription, and quantitative PCR

Total RNA was isolated from cultured cells using a phenol–chloroform extraction method. Briefly, cells were harvested, washed with phosphate-buffered saline (PBS), and lysed in TRIzol reagent. Following phase separation with chloroform, RNA was precipitated with isopropanol, washed with 75% ethanol, air-dried briefly, and resuspended in nuclease-free water. RNA concentration and purity were assessed spectrophotometrically.

Complementary DNA (cDNA) was synthesized from 1 µg of total RNA using random hexamer priming and reverse transcriptase according to standard protocols. Quantitative PCR (qPCR) was performed using SYBR Green chemistry with gene-specific primers (Supplementary Table 2). Relative gene expression was calculated using the ΔΔCt method with ACTB as the endogenous control. Melt-curve analysis was performed to confirm amplification specificity. All reactions were performed in technical triplicate, and experiments were repeated independently four times.

### Polysome profiling

To assess translational engagement, polysome profiling was performed using sucrose gradient ultracentrifugation. Cells were treated briefly with cycloheximide (100 µg ml⁻¹) to arrest translating ribosomes, washed in cycloheximide-containing PBS, and lysed in polysome lysis buffer containing Tris-HCl, KCl, MgCl₂, detergent, reducing agent, protease and phosphatase inhibitors, and cycloheximide. Lysates were clarified by sequential centrifugation at 4 °C.

Clarified lysates were layered onto linear 7–47% sucrose gradients prepared in low-salt buffer and subjected to ultracentrifugation to resolve ribonucleoprotein complexes. Following centrifugation, gradients were fractionated manually^51^, and fractions corresponding to free RNA, ribosomal subunits, monosomes, light polysomes, and heavy polysomes were collected based on relative gradient position. Polysome absorbance profiles were performed on cytoplasmic lysates layered on 7-47% sucrose gradients using a Biocomp Instruments Gradient Fractionation System (Tatamagouche, NS, Canada), with UV absorbance at 260 nm recorded using a Triax Flow Cell and associated software (Biocomp Instruments). RNA was isolated from individual polysome fractions using silica-based column purification, followed by ethanol precipitation to reconcentrate RNA. Purified RNA was resuspended in nuclease-free water and subjected to reverse transcription and quantitative PCR was performed as described above. To assess transcript levels within polysome fractions, relative transcript abundance was quantified as a fraction of total transcript across all collected fractions. Ct values from individual fractions were converted to linear expression values (2^−Ct), summed across all fractions for each transcript, and individual fraction values were expressed as a percentage of the total. This approach was used to assess transcript distribution across polysome fractions independent of housekeeping gene normalization.

### HLA Class I Immunoprecipitation

A549 cells were grown in indicated oxygen conditions for 48 hours. For experiments using 5-azacytiding (5AZA) or Decitabine (DAC), drugs were added at initial plating. To ensure adequate cell material, three 15cm tissue culture plate cultures were pooled per replicate. Cells were collected after careful washing with 1x PBS and lysed with immunoprecipitation lysis buffer (1% CHAPS (Millipore) 50mM Tris 150mM NaCl +1X HALT protease/phosphatase inhibitor (Thermo scientific) pH 8.0). Lysates were precleared of large insoluble aggregates and nucleic acids by centrifugation (4200-5000 RPM for 30 minutes). Protein A beads (Resyn Biosciences; protein A MAX) were coupled to 500ug w6/32 antibody in lysis buffer. Bead-antibody complexes were then washed 2x in lysis buffer and incubated with cleared lysate overnight at 4C with end over end rotation. Following incubation beads were washed 3x in DPBS and finally 2x in LC–MS/MS grade water. Washed beads were frozen at -80 and shipped to the Children’s Hospital of Philadelphia for elution and LC-MS/MS analysis.

### Mass spectrometry data acquisition

Samples underwent desalting using a C18 stage tip^52^ and were eluted with 28% acetonitrile/0.1% formic acid, followed by drying via vacuum centrifugation and reconstitution in 0.1% TFA/0.015% DDM containing iRT peptides. Randomized samples were then analyzed using a timsTOF Pro 2 mass spectrometer (Bruker Daltonics, Bremen, Germany) coupled with a nanoElute 2 via a captive spray ion source utilizing the Thunder-DDA-PASEF method^53^. Samples were injected onto a PepSep C18 column (25 cm × 150 µm × 1.5 µm, Bruker) using a linear gradient of 2–22% buffer B over 40 min, 28% B over 50 min, and 34% B over 60 min at a flow rate of 600 nL min⁻¹. Buffer A consisted of 0.1% formic acid (FA), while buffer B consisted of 0.1% FA in acetonitrile.

The captive spray source parameters were set to 1500 V, dry gas at 3.0 L min⁻¹, and a dry temperature of 180 °C. For data-dependent acquisition with PASEF (DDA-PASEF), settings included one full MS scan with a scan range of 100–1700 m/z. TIMS settings encompassed a 1/K₀ range of 0.60–1.60 V·s·cm⁻², ramp and accumulation times of 100 ms, and a ramp rate of 9.42 Hz. Mass spectra peak detection used an absolute threshold of 10, while mobility peak detection used an absolute threshold of 5,000. Each acquisition cycle comprised 10 PASEF scans with accumulation and ramp times of 100 ms each. Target intensity was set to 20,000 with an intensity threshold of 2,500, and dynamic exclusion was enabled with a duration of 0.4 min. Charge states 0–5 were permitted, and collision energy was set to 20 eV for 1/K₀ = 0.60 V·s·cm⁻² and 59 eV for 1/K₀ = 1.60 V·s·cm⁻².

### Mass spectrometry system suitability and quality control

Instrument suitability of the timsTOF Pro 2 system was monitored using QuiC software (Biognosys, Schlieren, Switzerland) based on analysis of spiked-in iRT peptides. In addition, as a quality control measure, a standard K562 (Promega) protein digest was injected before, midway through, and after the sample set using DDA mode. These quality control DDA data were analyzed using MaxQuant^54^, and results were visualized with the PTXQC package^55^ to track instrument performance.

### Mass spectrometry raw data processing

Raw mass spectrometry files were processed using MSFragger v21.1 within the FragPipe computational environment, with MSBooster enabled^20,21^. Searches were conducted against a human reference proteome from UniProt (Swiss-Prot and TrEMBL entries), supplemented with a list of 245 common protein contaminants. The default parameters of the Nonspecific-HLA workflow were used. Protein digestion was specified as nonspecific, and both precursor and fragment mass tolerances were set to 20 ppm. Variable modifications included protein N-terminal acetylation, oxidation of methionine, cysteinylation of cysteine, and pyroglutamate formation at N-terminal glutamine or glutamate residues. Peptide–spectrum matches were validated using Percolator with a minimum probability threshold of 0.5. MS1-level label-free quantification was performed using IonQuant^56^ without enabling match-between-runs.

### Immunopeptide post-search filtering and peptide compilation

Following database searching, peptide-spectrum matches were filtered to an experiment-wide false discovery rate (FDR) of 2% at the peptide level using a target–decoy strategy. For downstream analyses, peptide identifications were collapsed by amino acid sequence, irrespective of charge state or repeated spectral identification. To reduce the influence of stochastic sampling, only immunopeptides detected in at least two of three biological replicates within any experimental group were retained for quantitative and comparative analyses.

### Neoantigen binding prediction and analysis

To identify candidate neoantigens in A549 cells, the reference sequence database was augmented to include predicted HLA-binding immunopeptides harboring single nucleotide polymorphisms (SNPs) identified from publicly available whole-exome sequencing data for A549 cells^57,58^. For each variant, 25–amino acid sequences centered on the SNP were used to generate predicted HLA-binding peptides using NetMHCpan v4.0. Peptides containing nonsynonymous variants were evaluated at lengths of 8–12 amino acids against the corresponding HLA class I alleles expressed by A549 cells (HLA-A*25:01, HLA-A*30:01, HLA-B*18:01, HLA-B*44:03, HLA-C*12:03, and HLA-C*16:01). Both high- and low-affinity predicted binders were retained to avoid biasing candidate selection toward only the strongest predicted interactions. Binding predictions were used solely to prioritize candidate neoantigen peptides for inclusion in the augmented reference library and were not interpreted as evidence of peptide presentation.

Raw LC–MS/MS data were re-analyzed against this augmented database using the same search parameters as in the primary immunopeptidomics analysis, including fully nonspecific digestion and target–decoy-based validation. Peptide-spectrum matches were filtered using the same PSM-and peptide-level false discovery rate (FDR) thresholds applied in the primary analysis. To increase sensitivity for detection of low-abundance variant-derived peptides across biological replicates, match-between-runs (MBR) was enabled in IonQuant with a relaxed ion-level FDR threshold of 5%. In addition to directly identified peptide-spectrum matches, inferred peptide features supported by consistent retention time behavior, spectral similarity, and agreement with expected HLA-binding motifs were evaluated. Candidate neoantigens were interpreted as putative identifications and were required to be incompatible with any wild-type peptide explanation and to uniquely map to variant-derived sequences, consistent with common practice in exploratory immunopeptidomic analyses^59^.

### Peptide-to-protein mapping and peptide diversity analyses

Identified immunopeptides were mapped to their source proteins by exact sequence matching. For analyses of peptide diversity, each unique peptide sequence was counted once per sample. Diversity metrics were calculated per protein using the number of distinct immunopeptides derived from that protein within each experimental condition. Group-level peptide sets were generated by aggregating unique peptides across biological replicates within each experimental group.

### Anchor residue and motif analysis

To assess differences in HLA anchor residue usage across experimental conditions, peptides were grouped by inferred HLA restriction and peptide length. Amino acid frequencies at canonical anchor positions (including P2 and the C-terminal Ω residue for MHC class I peptides) were calculated for each condition. Frequency vectors were subjected to centered log-ratio (CLR) transformation to account for the compositional nature of residue frequency data.

Differences in anchor residue composition between conditions were evaluated using permutational multivariate analysis of variance (PERMANOVA) applied to CLR-transformed frequency data. Where indicated, post hoc analyses were performed to identify residues contributing most strongly to observed differences. Motif visualizations were generated using MHCviz and related tools.

### Generation of HIF1A, EPAS1 (HIF2A), and HIF1A/EPAS1 knockout cell lines

HIF1A, EPAS1 (HIF2A), and combined HIF1A/EPAS1 KO A549 cell lines were generated using CRISPR/Cas9 ribonucleoprotein (RNP)–mediated genome editing. CRISPR guide RNAs targeting human HIF1A OR EPAS1 were selected from predesigned Alt-R CRISPR-Cas9 guide sets (Integrated DNA Technologies, IDT). Two independent guide RNAs were used simultaneously for both HIF1A and EPAS1 to increase editing efficiency (Supplementary Table 3). Guide RNAs were duplexed with Alt-R trans-activating CRISPR RNA (tracrRNA; IDT) and complexed with recombinant Alt-R S.p. Cas9 nuclease V3 (IDT) according to the manufacturer’s instructions to form RNP complexes. Briefly, equimolar crRNA and tracrRNA were annealed and incubated with Cas9 protein at room temperature to generate RNPs immediately prior to electroporation.

A549 cells (1 × 10⁷) were washed, resuspended in Opti-MEM (Gibco), and combined with RNP complexes and Alt-R Cas9 Electroporation Enhancer (IDT). Cells were electroporated in 4-mm gap cuvettes using a square-wave electroporation system (single pulse, ∼225–250 V, 30 ms). Following electroporation, cells were rested in cuvettes for 30 min at 37 °C and then transferred to complete growth medium. Cell viability after electroporation was typically 25–40%. Single-cell clones were isolated by limiting dilution in 96-well plates and expanded over several weeks. HIF1A KO, EPAS1 KO, and HIF1A/EPAS1 double KO clones were identified by immunoblotting. For validation, cells were exposed to cobalt chloride (CoCl₂) to stabilize HIF proteins prior to lysis; loss of inducible HIF-1α and/or HIF-2α protein expression was confirmed by western blotting (Figure 3e). The HIF1A/EPAS1 double knockout line was generated sequentially by targeting EPAS1 in a previously validated HIF1A KO clone using the same CRISPR/Cas9 RNP approach. Multiple independent KO clones were screened, and representative clones with complete loss of inducible HIF protein expression were selected for downstream experiments.

### Nanopore direct RNA sequencing and m5C analysis

Total RNA was isolated from A549 cells preconditioned to normoxia (20% O₂) or hypoxia (2% O₂) for 48 h prior to IFN-γ stimulation (n = 3). Polyadenylated RNA was prepared according to the manufacturer’s recommendations and used as input for direct RNA library preparation using the SQK-RNA004 chemistry kit (Oxford Nanopore Technologies). Libraries were sequenced on PromethION flow cells to obtain native RNA reads without reverse transcription or amplification.

Raw nanopore signal data were base-called using Dorado v1.0.2 (Oxford Nanopore Technologies). Modified base detection was performed during base-calling using the ONT RNA004 SUP modified-base model (rna004_130bps_sup@v5.1.0), which supports prediction of cytosine-5 methylation (m5C) from direct RNA sequencing data. Reads were aligned to the human reference genome (GRCh38), and per-base modification probability estimates were extracted from aligned reads.

For downstream analyses, m5C calls were filtered to retain high-confidence sites, defined as a predicted methylation probability ≥ 0.8 and a minimum total read coverage of 50 reads per site. Predicted m5C calls were summarized across transcripts and compared between normoxic and hypoxic conditions. Transcriptome-wide differential analyses were performed to identify potential hypoxia-associated shifts in RNA methylation patterns, with analyses restricted to sites meeting coverage and confidence thresholds in all compared conditions.

For visualization and pathway-level analyses, differential m5C signals were aggregated at the gene level. Volcano plots were generated by plotting effect size (log₂ fold change in mean methylation probability) against –log₁₀ P values derived from gene-level statistical testing. For gene-level differential analyses, P values were corrected for multiple testing using the Benjamini–Hochberg false discovery rate (FDR) procedure, with q < 0.05 considered significant. Gene set enrichment analysis (GSEA) was performed using ranked gene lists based on signed differential m5C signal, and pathway-level significance was assessed using FDR correction as implemented in the GSEA framework, with q < 0.25 considered significant.

### m5C methylated RNA immunoprecipitation sequencing (MeRIP-seq)

m5C methylated RNA immunoprecipitation sequencing (MeRIP-seq) was performed to assess transcriptome-wide cytosine-5 methylation patterns. A549 cells were cultured under the indicated experimental conditions, and total RNA (approximately 100–400 µg per condition) was isolated using TRIzol reagent according to the manufacturer’s protocol. RNA integrity was assessed prior to downstream processing. Total RNA was then chemically fragmented to an average length of ∼100 nucleotides using RNA fragmentation buffer (100 mM Tris-HCl, pH 7.0; 100 mM ZnCl₂) by incubation at 95 °C for 5 min. Fragmentation reactions were quenched by addition of 0.5 M EDTA, and fragment size distributions were confirmed using capillary electrophoresis. Fragmented RNA was purified by ethanol precipitation using sodium acetate and glycogen as carrier and resuspended in RNase-free water.

For immunoprecipitation, fragmented RNA was incubated in immunoprecipitation (IP) buffer (750 mM NaCl, 0.5% IGEPAL CA-630, 50 mM Tris-HCl, pH 7.4), supplemented with RNase inhibitor, anti-m5C antibody (Diagenode, 33D3, 1 ug/mL), and Protein G magnetic beads. Reactions were incubated with end-over-end rotation for 2 h at 4 °C. Beads were washed three times with IP buffer to remove unbound RNA. Bound RNA was eluted by incubation with elution buffer containing free m5C nucleoside and RNase inhibitor for 1 h at 4 °C with agitation. Eluted RNA was recovered by ethanol precipitation, washed with 75% ethanol, air-dried, and resuspended in RNase-free water. Matched input RNA samples were processed in parallel. Immunoprecipitated and input RNA fractions were converted to cDNA libraries using standard protocols and submitted for high-throughput sequencing.

### MeRIP-seq peak calling and differential analysis

m5C-enriched regions were identified and quantified using the exomePeak2 package (Bioconductor). Sequencing reads from immunoprecipitated (IP) and matched input RNA samples were analyzed using a negative binomial modeling framework to identify regions of enrichment relative to input. Analyses were performed using the human reference genome (GRCh38) and corresponding gene annotation (GENCODE). Peak calling and differential enrichment analyses were conducted using exomePeak2 with recommended default parameters, including GC content modeling. Biological replicates were incorporated into the statistical model where applicable. Differential m5C enrichment between normoxic (20% O₂) and hypoxic (2% O₂) conditions was assessed at the peak level. Resulting P values were adjusted for multiple testing using the Benjamini–Hochberg false discovery rate (FDR) procedure. For transcriptome-wide interpretation, peak-level signals were additionally aggregated at the transcript and gene levels. No reproducible hypoxia-associated differential m5C enrichment was detected after multiple-testing correction.

### Immunofluorescence staining and RNA fluorescence in situ hybridization

For cell-based immunofluorescence and RNA fluorescence in situ hybridization (RNA FISH), A549 cells were seeded on 18-mm glass coverslips and cultured under normoxic (20% O₂) or hypoxic (2% O₂) conditions as indicated. Cells were treated with IFN-γ for 24 h, with 5-azacytidine or decitabine added at the time of plating where indicated. Sodium arsenite treatment (500 µM for 2 h) was used as a positive control for stress granule formation. Cells were washed with phosphate-buffered saline (PBS) and fixed in 4% paraformaldehyde for 15 min at room temperature. For immunofluorescence, cells were permeabilized with ice-cold methanol, blocked, and incubated with anti-G3BP1, and anti-m5C-FITC conjugated primary antibodies. G3BP1 was detected using an Alexa Fluor 594–conjugated secondary antibody, and nuclei were counterstained with DAPI. m5C signal was acquired as part of the imaging panel and is included in the raw image data.

For RNA FISH, cells were processed according to the Stellaris™ RNA FISH protocol for adherent cells with minor modifications. Following fixation and permeabilization, cells were hybridized overnight at 37 °C with probe-containing hybridization buffer targeting PSMB8, PSMB9, or PSMB10 transcripts conjugated to Quasar 670 dye. Probes were designed using the custom Stellaris^TM^ Probe Designer (Biosearch Technologies) (Supplementary Table 4). After post-hybridization washes, nuclei were counterstained with DAPI. Where combined RNA FISH and immunofluorescence was performed, immunostaining was carried out prior to final mounting.

Formalin-fixed, paraffin-embedded (FFPE) human non-small cell lung cancer tissue sections were obtained through the University of Iowa Tissue Procurement and BioMER Core Facility. Tissue sections were derived stage II–III NSCLC tumors. FFPE sections were deparaffinized in xylene and rehydrated through graded ethanol to water. Antigen retrieval was performed in Tris-EDTA buffer containing detergent by heat-induced epitope retrieval, followed by cooling to room temperature. Sections were permeabilized, blocked, and incubated with primary antibodies overnight at 4 °C. IP subunits were detected using Alexa Fluor 647–conjugated antibodies, CA9 was detected using an Alexa Fluor 594–conjugated secondary antibody, and nuclei were counterstained with DAPI. Sections were mounted in antifade mounting medium prior to imaging.

### Confocal image acquisition and processing

Fluorescence imaging was performed using a laser scanning confocal microscope equipped with a 63× oil immersion objective. Images were acquired as single optical sections without z-stacking. Fluorescence channels were acquired sequentially to minimize spectral bleed-through. All images were processed using identical, linear adjustments to brightness and contrast applied uniformly across all experimental conditions within each experiment. No nonlinear image processing or deconvolution was applied.

### Superpixel-based regional quantification and correlation analysis

Multiplex fluorescence images were analyzed in QuPath. Tumor regions were delineated as annotations to restrict quantification to tissue compartments of interest (ROIs). Within each ROI, images were partitioned into superpixels using the Simple Linear Iterative Clustering (SLIC) algorithm. For each superpixel, per-channel mean fluorescence intensity was computed for the CA9 (hypoxia) channel and for the relevant IP catalytic subunit. Superpixels were then assigned to one of four quadrants based on CA9 and IP catalytic subunit intensities (double negative, CA9+ alone, IP subunit+ alone, or co-regional). Thresholds defining quadrant boundaries were determined using an objective intensity cutoff approach consistent with established guidance for fluorescence thresholding and background exclusion^60^. In addition, Spearman rank correlations (ρ) were calculated between CA9 and PSMB superpixel mean intensities within each tumor ROI. Only samples with ≥1% CA9-positive area were included in correlation summaries (7 of 9 tumors), to avoid unstable estimates from slides lacking measurable hypoxic signal.

### Statistical analysis

Statistical analyses were performed using GraphPad Prism or R. Unless otherwise stated, data are presented as mean ± SEM. Statistical tests, exact n values, and definitions of replicates are specified in the corresponding figure legends. Two-tailed Student’s t-tests were used for pairwise comparisons unless otherwise indicated. For analyses involving multiple comparisons, appropriate corrections were applied as described in the relevant Methods sections or figure legends. P ≤ 0.05 was considered statistically significant.

## Data Availability

Raw and processed immunopeptidomics mass spectrometry data generated in this study have been deposited in the ProteomeXchange Consortium via the PRIDE partner repository under accession number PXD074571. Nanopore direct RNA sequencing data have been deposited in the NCBI Sequence Read Archive (SRA) under BioProject accession number (accession number pending). m5C methylated RNA immunoprecipitation sequencing (MeRIP-seq) data have been deposited in the Gene Expression Omnibus (GEO) under accession number (accession number pending). All other data supporting the findings of this study are available in the Article, its Extended Data, and its Supplementary Information. Raw data are available from the corresponding author upon request.

## Acknowledgements

This work was supported by the NIH grant CA214285, and the American Cancer Society grant CAT-24-1414241-01-CAT. Flow cytometry data was acquired in the University of Iowa’s Flow Cytometry Facility, which is a Carver College of Medicine / Holden Comprehensive Cancer Center core research facility. Microscopy data was acquired in the University of Iowa Central Microscopy Research Facility. Nanopore and m5C MeRIP-seq data were obtained at the Genomics Division of the Iowa Institute of Human Genetics (RRID: SCR_023422) which is supported, in part, by the University of Iowa Carver College of Medicine and the Holden Comprehensive Cancer Center (National Cancer Institute of the National Institutes of Health under Award Number P30CA086862). Proteomics was performed by the Children’s Hospital of Philadelphia (CHOP), Penn Medicine Hospital of The University of Pennsylvania, Proteomics Core Facility.

## Author Contributions

All authors made significant contributions to the manuscript. A.W.M. conceived of the study.A.W.M., M.G.S., and A.R.R. contributed to the design of the project and experiments. A.W.M., A.W.M., M.G.S., A.R.R, and H.P. set up experiments and performed cell culture work, western blotting, RNA isolation, protein isolation, immunoprecipitation, and limiting dilution for clone derivation. A.W.M. performed CRISPER/Cas9. M.G.S. set up experiments and prepared resulting samples for immunopeptidomics analysis. M.G.S. performed and analyzed flow cytometry. A.W.M. performed anchor residue analysis. A.W.M. and M.G.S. performed neoantigen analysis and analysis of immunopeptide results. L.S. and H.F. performed mass spectrometry and mass spectrometry analysis. M.G.S, A.R.R., and H.P. set up experiments and prepared resulting samples for nanopore sequencing and m5C MeRIP-seq. A.W.M. performed nanopore m5C base-calling and analysis and MeRIP-seq analysis. A.R.R. set up experiments for and performed all RT-qPCR and polysome profiling experiments. M.G.S. and H.P. set up experiments for and performed stress granule staining and FISH. A.W.M. performed confocal imaging acquisition and imaging analysis. L.B.M.Jr. performed polysome profiling and provided resulting traces. N.C. set up experiments for and performed western blotting for CoCl_2_-treatment experiments. C.G. aided in RT-qPCR and set up and performed m5C-realted assays. A.W.M., A.R.R., M.G.S., and H.P. wrote the manuscript.

## Main References

1 Ayers, M. et al. IFN-gamma-related mRNA profile predicts clinical response to PD-1 blockade. J Clin Invest 127, 2930–2940 (2017). 10.1172/JCI91190

2 Thompson, J. C. et al. Gene signature of antigen processing and presentation machinery predicts response to checkpoint blockade in non-small cell lung cancer (NSCLC) and melanoma. J Immunother Cancer 8 (2020). 10.1136/jitc-2020-000974

3 Grasso, C. S. et al. Conserved Interferon-gamma Signaling Drives Clinical Response to Immune Checkpoint Blockade Therapy in Melanoma. Cancer Cell 38, 500–515 e503 (2020). 10.1016/j.ccell.2020.08.005

4 Kalaora, S. et al. Immunoproteasome expression is associated with better prognosis and response to checkpoint therapies in melanoma. Nat Commun 11, 896 (2020). 10.1038/s41467-020-14639-9

5 Nielsen, M. et al. Coexisting Alterations of MHC Class I Antigen Presentation and IFNgamma Signaling Mediate Acquired Resistance of Melanoma to Post-PD-1 Immunotherapy. Cancer Immunol Res 10, 1254–1262 (2022). 10.1158/2326-6066.CIR-22-0326

6 Tripathi, S. C. et al. Immunoproteasome deficiency is a feature of non-small cell lung cancer with a mesenchymal phenotype and is associated with a poor outcome. Proc Natl Acad Sci U S A 113, E1555–1564 (2016). 10.1073/pnas.1521812113

7 Guillaume, B. et al. Analysis of the processing of seven human tumor antigens by intermediate proteasomes. J Immunol 189, 3538–3547 (2012). 10.4049/jimmunol.1103213

8 Morel, S. et al. Processing of some antigens by the standard proteasome but not by the immunoproteasome results in poor presentation by dendritic cells. Immunity 12, 107–117 (2000). 10.1016/s1074-7613(00)80163-6

9 Rana, P. S. et al. Immunoproteasome Activation Expands the MHC Class I Immunopeptidome, Unmasks Neoantigens, and Enhances T-cell Anti-Myeloma Activity. Mol Cancer Ther 23, 1743–1760 (2024). 10.1158/1535-7163.MCT-23-0931

10 Garcia-Lora, A., Algarra, I. & Garrido, F. MHC class I antigens, immune surveillance, and tumor immune escape. J Cell Physiol 195, 346–355 (2003). 10.1002/jcp.10290

11 Neefjes, J., Jongsma, M. L., Paul, P. & Bakke, O. Towards a systems understanding of MHC class I and MHC class II antigen presentation. Nat Rev Immunol 11, 823–836 (2011). 10.1038/nri3084

12 Wang, S., He, Z., Wang, X., Li, H. & Liu, X. S. Antigen presentation and tumor immunogenicity in cancer immunotherapy response prediction. Elife 8 (2019). 10.7554/eLife.49020

13 Gaczynska, M., Rock, K. L., Spies, T. & Goldberg, A. L. Peptidase activities of proteasomes are differentially regulated by the major histocompatibility complex-encoded genes for LMP2 and LMP7. Proc Natl Acad Sci U S A 91, 9213–9217 (1994). 10.1073/pnas.91.20.9213

14 Toes, R. E. et al. Discrete cleavage motifs of constitutive and immunoproteasomes revealed by quantitative analysis of cleavage products. J Exp Med 194, 1–12 (2001). 10.1084/jem.194.1.1

15 Boegel, S. et al. HLA and proteasome expression body map. BMC Med Genomics 11, 36 (2018). 10.1186/s12920-018-0354-x

16 Yamamoto, K. et al. Autophagy promotes immune evasion of pancreatic cancer by degrading MHC-I. Nature 581, 100–105 (2020). 10.1038/s41586-020-2229-5

17 Estephan, H. et al. Hypoxia promotes tumor immune evasion by suppressing MHC-I expression and antigen presentation. EMBO J 44, 903–922 (2025). 10.1038/s44318-024-00319-7

18 Byrne, M. B., Leslie, M. T., Gaskins, H. R. & Kenis, P. J. A. Methods to study the tumor microenvironment under controlled oxygen conditions. Trends Biotechnol 32, 556–563 (2014). 10.1016/j.tibtech.2014.09.006

19 Cui, Y. L., Wang, X. & Li, X. F. (18)F-fluoromisonidazole PET reveals spatial and temporal heterogeneity of hypoxia in mouse models of human non-small-cell lung cancer. Future Oncol 11, 2841–2849 (2015). 10.2217/fon.15.205

20 Kong, A. T., Leprevost, F. V., Avtonomov, D. M., Mellacheruvu, D. & Nesvizhskii, A. I. MSFragger: ultrafast and comprehensive peptide identification in mass spectrometry-based proteomics. Nat Methods 14, 513–520 (2017). 10.1038/nmeth.4256

21 Yang, K. L. et al. MSBooster: improving peptide identification rates using deep learning-based features. Nat Commun 14, 4539 (2023). 10.1038/s41467-023-40129-9

22 Diesinger, I. et al. Toward a more complete recognition of immunoreactive antigens in squamous cell lung carcinoma. Int J Cancer 102, 372–378 (2002). 10.1002/ijc.10714

23 Ito, S. et al. Expression of MAGE-D4, a novel MAGE family antigen, is correlated with tumor-cell proliferation of non-small cell lung cancer. Lung Cancer 51, 79–88 (2006). 10.1016/j.lungcan.2005.08.012

24 Palata, O. et al. Detection of tumor antigens and tumor-antigen specific T cells in NSCLC patients: Correlation of the quality of T cell responses with NSCLC subtype. Immunol Lett 219, 46–53 (2020). 10.1016/j.imlet.2020.01.001

25 Schag, K. et al. Identification of C-met oncogene as a broadly expressed tumor-associated antigen recognized by cytotoxic T-lymphocytes. Clin Cancer Res 10, 3658–3666 (2004). 10.1158/1078-0432.CCR-03-0640

26 Shomura, H. et al. Identification of epidermal growth factor receptor-derived peptides immunogenic for HLA-A2(+) cancer patients. Br J Cancer 90, 1563–1571 (2004). 10.1038/sj.bjc.6601728

27 Tsai, J. R. et al. Differential expression profile of MAGE family in non-small-cell lung cancer. Lung Cancer 56, 185–192 (2007). 10.1016/j.lungcan.2006.12.004

28 Yasumoto, K., Hanagiri, T. & Takenoyama, M. Lung cancer-associated tumor antigens and the present status of immunotherapy against non-small-cell lung cancer. Gen Thorac Cardiovasc Surg 57, 449–457 (2009). 10.1007/s11748-008-0433-6

29 Zhang, Y., Yang, D., Weng, L. & Wang, L. Early lung cancer diagnosis by biosensors. Int J Mol Sci 14, 15479–15509 (2013). 10.3390/ijms140815479

30 Ruppert, J. et al. Prominent role of secondary anchor residues in peptide binding to HLA-A2.1 molecules. Cell 74, 929–937 (1993). 10.1016/0092-8674(93)90472-3

31 Ivashkiv, L. B. IFNgamma: signalling, epigenetics and roles in immunity, metabolism, disease and cancer immunotherapy. Nat Rev Immunol 18, 545–558 (2018). 10.1038/s41577-018-0029-z

32 Hallermalm, K. et al. Tumor necrosis factor-alpha induces coordinated changes in major histocompatibility class I presentation pathway, resulting in increased stability of class I complexes at the cell surface. Blood 98, 1108–1115 (2001). 10.1182/blood.v98.4.1108

33 Ivashkiv, L. B. & Donlin, L. T. Regulation of type I interferon responses. Nat Rev Immunol 14, 36–49 (2014). 10.1038/nri3581

34 Semenza, G. L. Hypoxia-inducible factors in physiology and medicine. Cell 148, 399–408 (2012). 10.1016/j.cell.2012.01.021

35 Momparler, R. L. A Perspective on the Comparative Antileukemic Activity of 5-Aza-2’-deoxycytidine (Decitabine) and 5-Azacytidine (Vidaza). Pharmaceuticals (Basel) 5, 875–881 (2012). 10.3390/ph5080875

36 Adair, S. J. & Hogan, K. T. Treatment of ovarian cancer cell lines with 5-aza-2’-deoxycytidine upregulates the expression of cancer-testis antigens and class I major histocompatibility complex-encoded molecules. Cancer Immunol Immunother 58, 589–601 (2009). 10.1007/s00262-008-0582-6

37 Simova, J. et al. Immunotherapy augments the effect of 5-azacytidine on HPV16-associated tumours with different MHC class I-expression status. Br J Cancer 105, 1533–1541 (2011). 10.1038/bjc.2011.428

38 Munteanu, R. A. et al. 5-Azacytidine treatment inhibits the development of lung cancer models via epigenetic reprogramming and activation of cellular pathways with anti-tumor activity. Med Pharm Rep 97, 488–506 (2024). 10.15386/mpr-2777

39 Liu, L., Wise, D. R., Diehl, J. A. & Simon, M. C. Hypoxic reactive oxygen species regulate the integrated stress response and cell survival. J Biol Chem 283, 31153–31162 (2008). 10.1074/jbc.M805056200

40 Attwood, K. M. et al. Raloxifene prevents stress granule dissolution, impairs translational control and promotes cell death during hypoxia in glioblastoma cells. Cell Death Dis 11, 989 (2020). 10.1038/s41419-020-03159-5

41 Tourriere, H. et al. The RasGAP-associated endoribonuclease G3BP assembles stress granules. J Cell Biol 160, 823–831 (2003). 10.1083/jcb.200212128

42 Blankenstein, T., Coulie, P. G., Gilboa, E. & Jaffee, E. M. The determinants of tumour immunogenicity. Nat Rev Cancer 12, 307–313 (2012). 10.1038/nrc3246

43 Chowell, D. et al. Patient HLA class I genotype influences cancer response to checkpoint blockade immunotherapy. Science 359, 582–587 (2018). 10.1126/science.aao4572

44 Jhunjhunwala, S., Hammer, C. & Delamarre, L. Antigen presentation in cancer: insights into tumour immunogenicity and immune evasion. Nat Rev Cancer 21, 298–312 (2021). 10.1038/s41568-021-00339-z

45 Sade-Feldman, M. et al. Resistance to checkpoint blockade therapy through inactivation of antigen presentation. Nat Commun 8, 1136 (2017). 10.1038/s41467-017-01062-w

46 Cardenas-Navia, L. I. et al. The pervasive presence of fluctuating oxygenation in tumors. Cancer Res 68, 5812–5819 (2008). 10.1158/0008-5472.CAN-07-6387

47 Bruni, D., Angell, H. K. & Galon, J. The immune contexture and Immunoscore in cancer prognosis and therapeutic efficacy. Nat Rev Cancer 20, 662–680 (2020). 10.1038/s41568-020-0285-7

48 Thiruthaneeswaran, N. et al. Lost in application: Measuring hypoxia for radiotherapy optimisation. Eur J Cancer 148, 260–276 (2021). 10.1016/j.ejca.2021.01.039

49 Ivanov, P., Kedersha, N. & Anderson, P. Stress Granules and Processing Bodies in Translational Control. Cold Spring Harb Perspect Biol 11 (2019). 10.1101/cshperspect.a032813

50 Bhuvanagiri, M. et al. 5-azacytidine inhibits nonsense-mediated decay in a MYC-dependent fashion. EMBO Mol Med 6, 1593–1609 (2014). 10.15252/emmm.201404461

## Method References

51 Sobhany, M. & Stanley, R. E. Polysome Profiling without Gradient Makers or Fractionation Systems. J Vis Exp (2021). 10.3791/62680

52 Rappsilber, J., Mann, M. & Ishihama, Y. Protocol for micro-purification, enrichment, pre-fractionation and storage of peptides for proteomics using StageTips. Nat Protoc 2, 1896–1906 (2007). 10.1038/nprot.2007.261

53 Gomez-Zepeda, D. et al. Thunder-DDA-PASEF enables high-coverage immunopeptidomics and is boosted by MS(2)Rescore with MS(2)PIP timsTOF fragmentation prediction model. Nat Commun 15, 2288 (2024). 10.1038/s41467-024-46380-y

54 Tyanova, S., Temu, T. & Cox, J. The MaxQuant computational platform for mass spectrometry-based shotgun proteomics. Nat Protoc 11, 2301–2319 (2016). 10.1038/nprot.2016.136

55 Bielow, C., Mastrobuoni, G. & Kempa, S. Proteomics Quality Control: Quality Control Software for MaxQuant Results. J Proteome Res 15, 777–787 (2016). 10.1021/acs.jproteome.5b00780

56 Yu, F., Haynes, S. E. & Nesvizhskii, A. I. IonQuant Enables Accurate and Sensitive Label-Free Quantification With FDR-Controlled Match-Between-Runs. Mol Cell Proteomics 20, 100077 (2021). 10.1016/j.mcpro.2021.100077

57 DepMap, Broad (2025). DepMap Public 24Q4. Dataset. depmap.org

58 Arafeh, R., Shibue, T., Dempster, J. M., Hahn, W. C. & Vazquez, F. The present and future of the Cancer Dependency Map. Nat Rev Cancer 25, 59–73 (2025). 10.1038/s41568-024-00763-x

59 Scull, K. E., Pandey, K., Ramarathinam, S. H. & Purcell, A. W. Immunopeptidogenomics: Harnessing RNA-Seq to Illuminate the Dark Immunopeptidome. Mol Cell Proteomics 20, 100143 (2021). 10.1016/j.mcpro.2021.100143

60 Costes, S. V. et al. Automatic and quantitative measurement of protein-protein colocalization in live cells. Biophys J 86, 3993–4003 (2004). 10.1529/biophysj.103.038422

