## Supplementary material for "Hypoxic stress granules trigger immunogenic dormancy in lung cancer": All extended data figures, legends, and supplementary data

### Extended Figures, Tables, and Extended Figure Legends

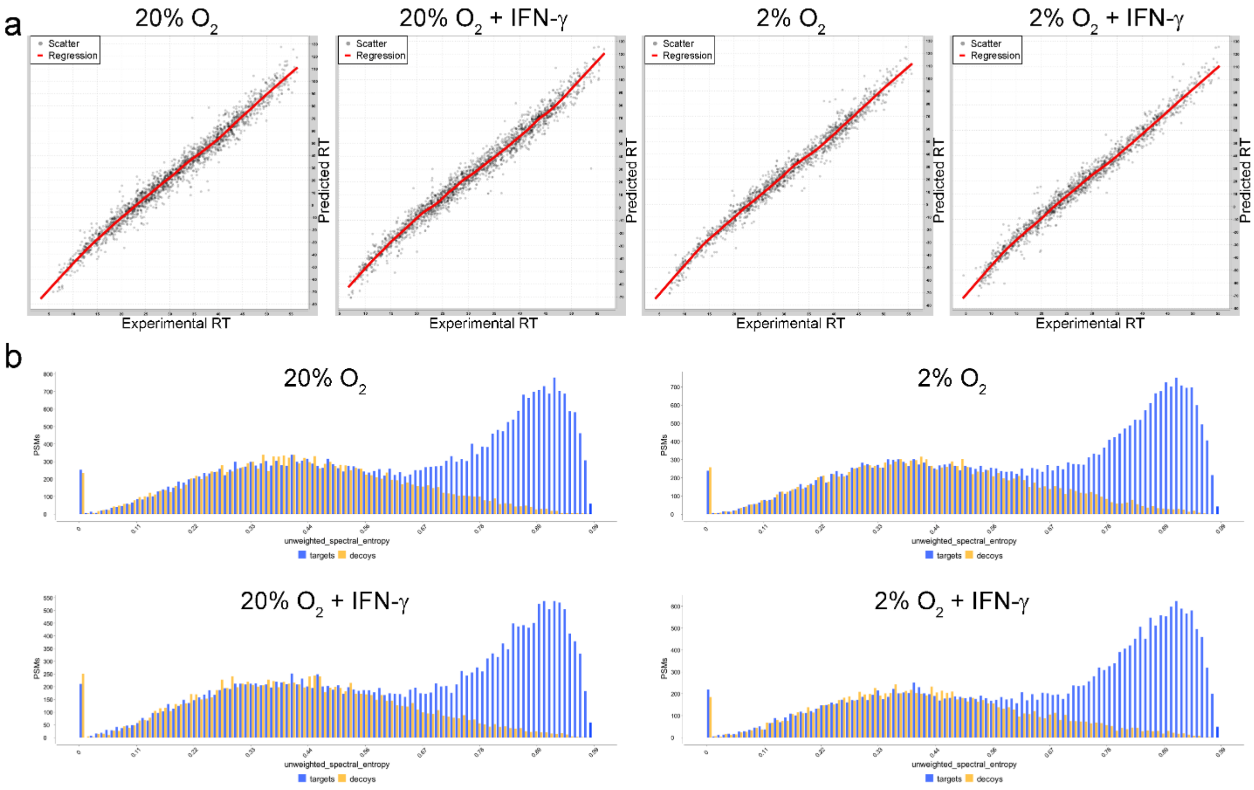


**Extended Data Figure 1: Quality control metrics for immunopeptidomics data processing**

**a**, Immunopeptidomic analysis was performed on HLA-immunoprecipitated eluates from A549 cells preconditioned in 20% $O_2$ or 2% $O_2$ for 48 h, followed by treatment with or without 10 ng/mL IFN-γ for an additional 24 h. MSBooster retention time (RT) calibration performance across representative samples are shown from each condition (20% $O_2$, 20% $O_2$ + IFN-γ, 2% $O_2$, and 2% $O_2$ + IFN-γ). Predicted RT values are plotted against experimental RT for confidently identified peptide–spectrum matches (PSMs). Red lines indicate LOESS regression fits, demonstrating strong concordance between predicted and observed RT across runs. **b**, Target–decoy score distributions for each condition based on unweighted spectral entropy after rescoring. Histograms display the separation of target (blue) and decoy (orange) PSMs, illustrating effective discrimination and controlled false discovery rate across all samples.

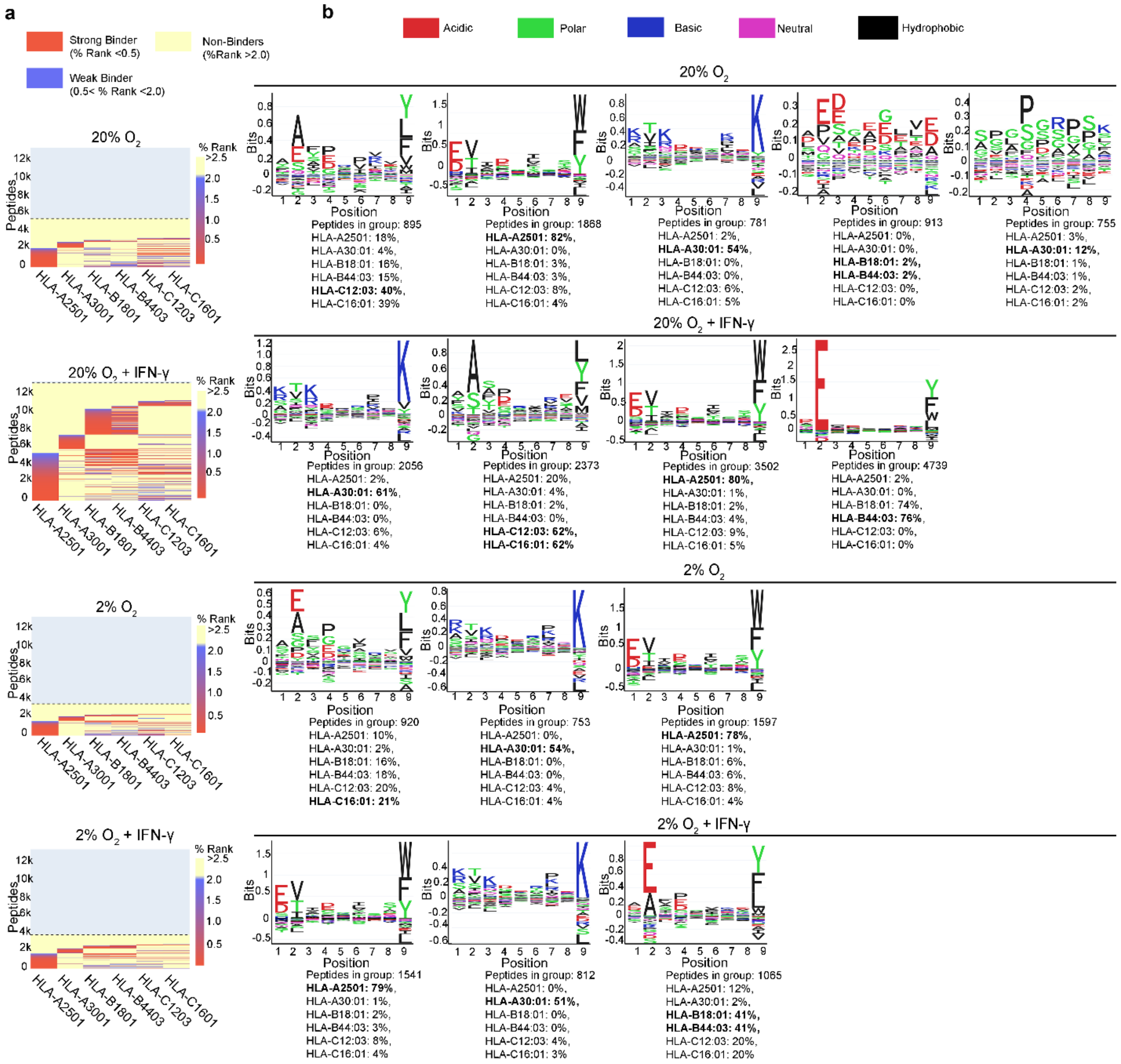


**Extended Data Figure 2: Unsupervised motif visualization of MHC class I–associated immunopeptides. a-b**, HLA class I–associated peptides were profiled from A549 cells cultured for 48 h under normoxia (20% $O_2$) or hypoxia (2% $O_2$) followed by 24 h stimulation in the presence or absence of IFN-γ. HLA class I–bound peptides were isolated by HLA immunoprecipitation, identified by LC–MS/MS, and analyzed using the MHCVizPipe immunopeptidomics workflow.

**a**, Heatmaps display NetMHCpan-predicted binding of individual peptides to expressed HLA class I alleles. Peptides were classified as strong binders (%Rank < 0.5) or weak binders (0.5 ≤ %Rank < 2.0).**b**, Unsupervised Gibbs clustering was performed on eluted peptide sequences within each condition to identify peptide-binding motif clusters. Sequence logos depict position-specific amino acid frequencies within each cluster. Total column height represents information content (bits), and letter height indicates relative amino acid contribution. Amino acid colors denote biochemical class.

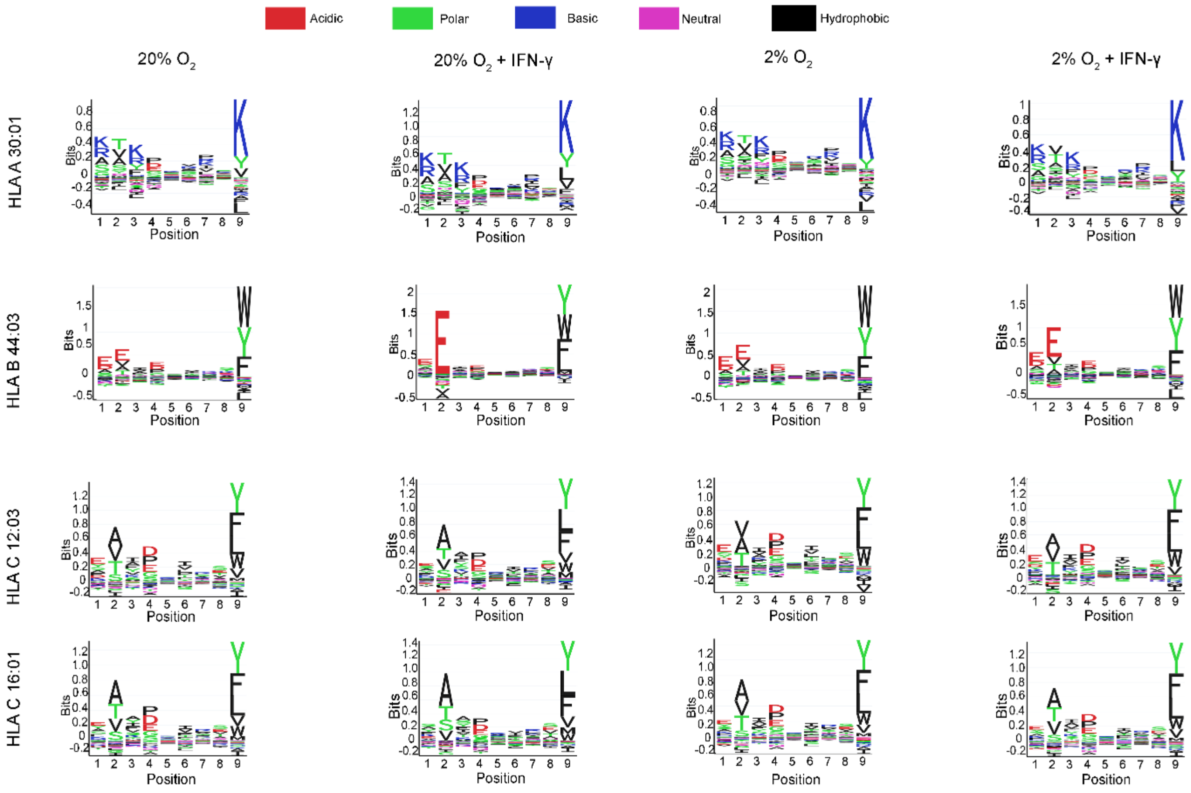


**Extended Data Figure 3: HLA allele specific motif visualization of MHC class I-associated immunopeptides**

Immunopeptidomics workflow and peptide identification were performed as described in Extended Data Fig. 2. Gibbs clustering was used to derive allele-specific peptide-binding motifs from eluted peptides. Sequence logos are organized by HLA allele (rows) and experimental condition (columns).

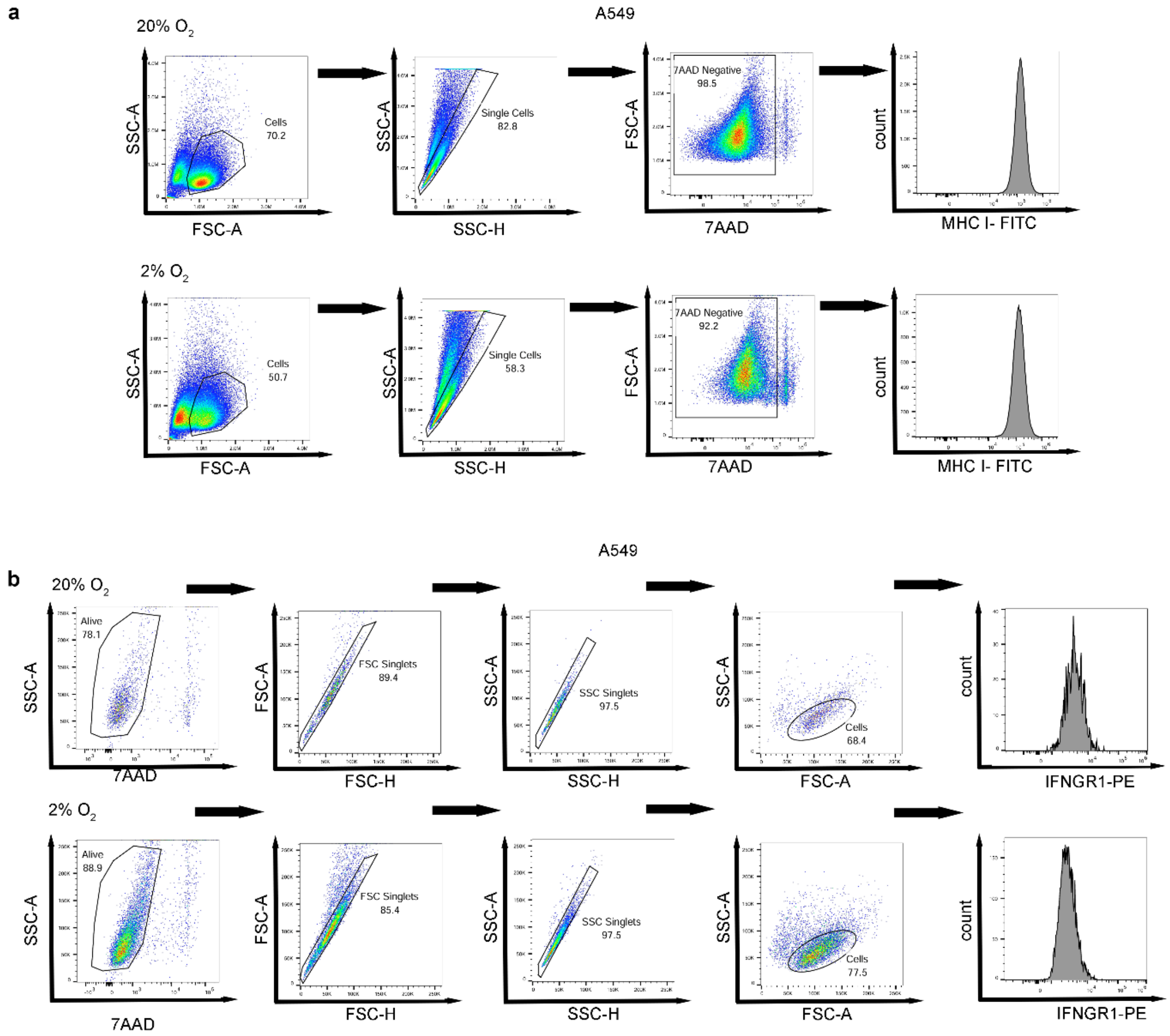


**Extended Data Figure 4: Flow cytometry gating strategy.**

**a-b**, Flow cytometry gating strategy used for analysis of cell surface protein expression in FlowJo. **a**, Sequential gating strategy for quantification of MHC class I surface expression on A549 cells cultured under normoxia or hypoxia for 48 h followed by 24 h IFN-γ stimulation. Viable cells were identified by exclusion of 7-AAD–positive events. **b**, Sequential gating strategy for quantification of IFNGR1 surface expression under the same experimental conditions. Non-viable cells were excluded using 7-AAD.

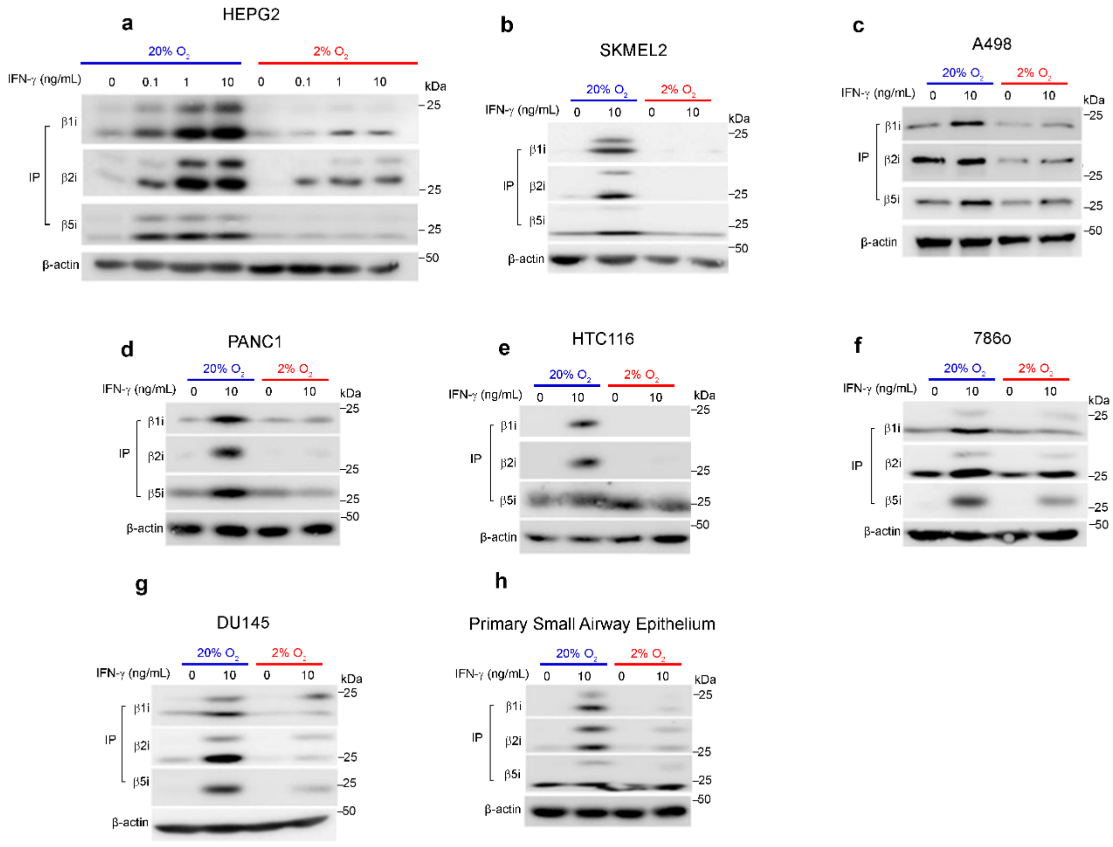


**Extended Data Figure 5: Analysis of inducible immunoproteasome expression across cancer cell lines**

**a-h,** Immunoblot analysis of lysates from Human hepatocellular carcinoma HepG2 (**a**); human melanoma SKMEL2 (**b**), human renal cell carcinoma A498 (**c**), human pancreatic cancer PANC1 (**d**), human colorectal cancer HTC116 (**e**), human renal cell carcinoma 786o (**f**), human prostate cancer DU145 (**g**), and cadaver donor primary airway epithelium (**h)** were cultured in indicated oxygen conditions for 48 hours followed by 24 hours of IFN-γ stimulation prior to cell lysate analysis by immunoblot for expression of inducible immunoproteasome subunits β1i, β2i, β5i.  β-actin included as a loading control for **a-h**.

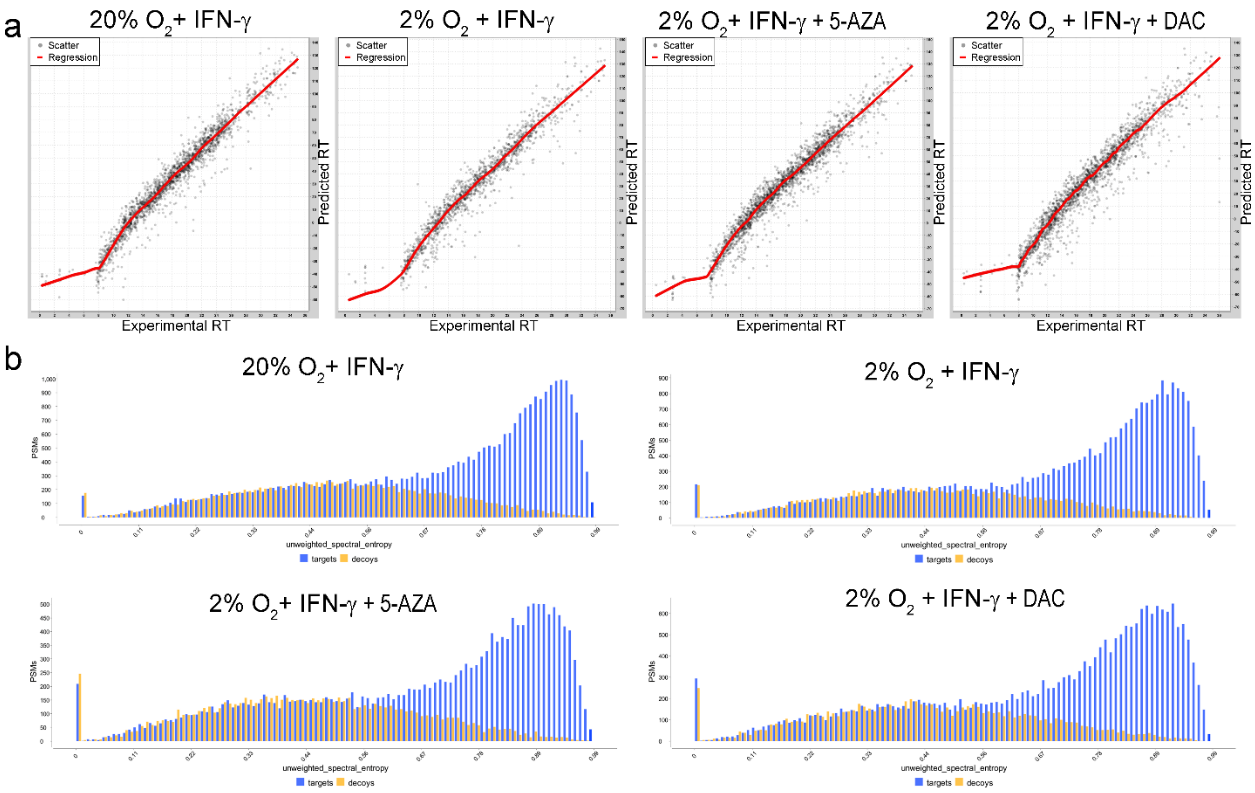


**Extended Data Figure 6: Quality control metrics for immunopeptidomics data processing with 5-AZA or DAC treatment**

**a**, Immunopeptidomic analysis was performed on HLA-immunoprecipitated eluates from A549 cells preconditioned to 20% $O_2$ or 2% $O_2$ for 48 h with or without 5-AZA or DAC followed by treatment with the indicated dose of IFN-γ for an additional 24 h. MSBooster retention time (RT) calibration performance across representative samples are shown from each condition (20% $O_2$, 20% $O_2$ + IFN-γ, 2% $O_2$, and 2% $O_2$ + IFN-γ). Predicted RT values are plotted against experimental RT for confidently identified peptide–spectrum matches (PSMs). Red lines indicate LOESS regression fits, demonstrating strong concordance between predicted and observed RT across runs. **b**, Target–decoy score distributions for each condition based on unweighted spectral entropy after rescoring. Histograms display the separation of target (blue) and decoy (orange) PSMs, illustrating effective discrimination and controlled false discovery rate across all samples.

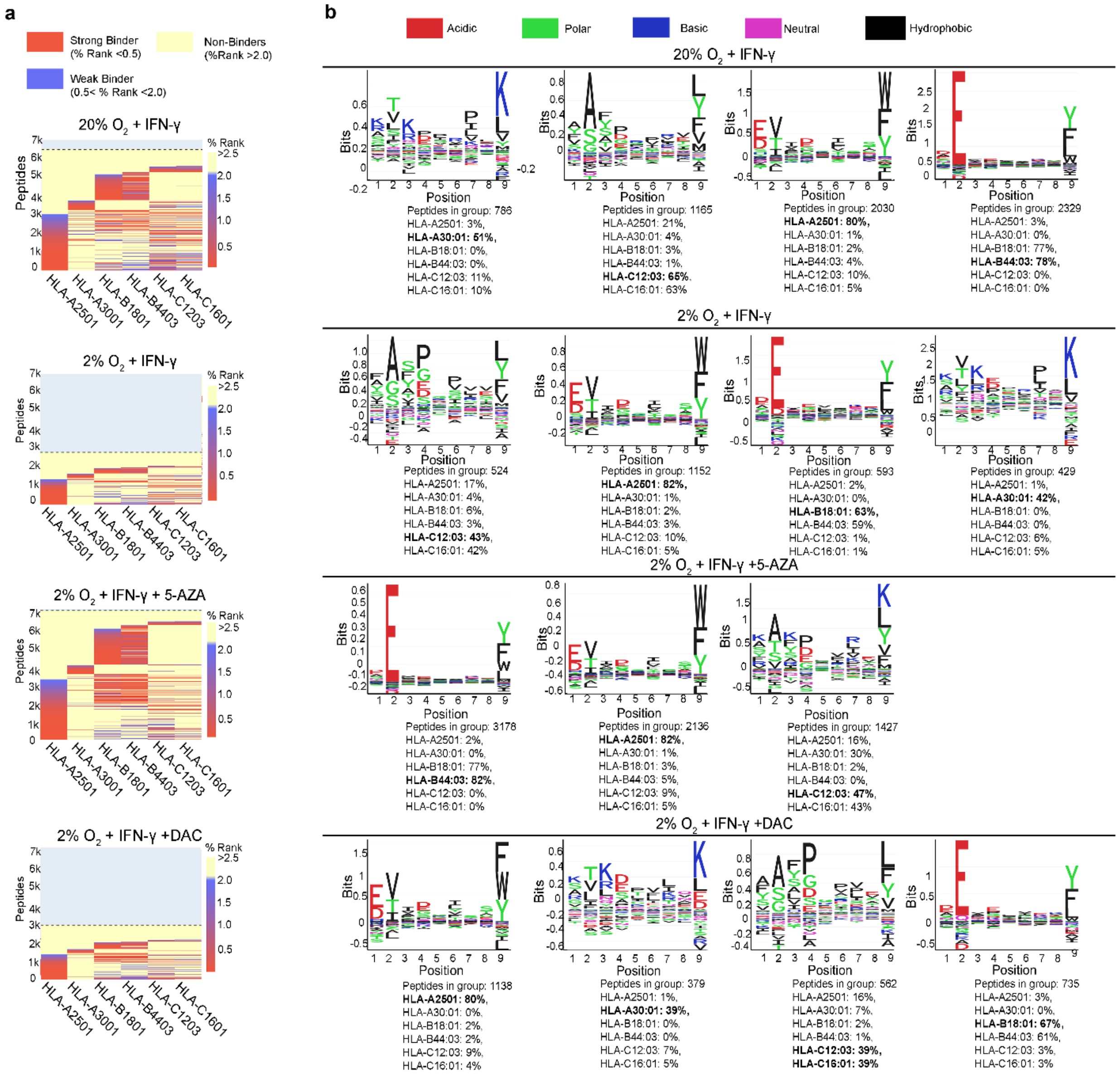


**Extended Data Figure 7: Unsupervised motif visualization of MHC class I–associated immunopeptides following epigenetic pretreatment.**

**a-b**, Immunopeptidomics workflow and peptide identification were performed as described in Extended Data Fig. 2. A549 cells were cultured for 48 h under normoxia or hypoxia in the presence or absence of 5-azacytidine (5-AZA) or decitabine (DAC), followed by 24 h stimulation with or without IFN-γ. **a**, Heatmaps display NetMHCpan-predicted binding of individual peptides to expressed HLA class I alleles. **b**, Unsupervised Gibbs clustering identified peptide-binding motif clusters, visualized as sequence logos.

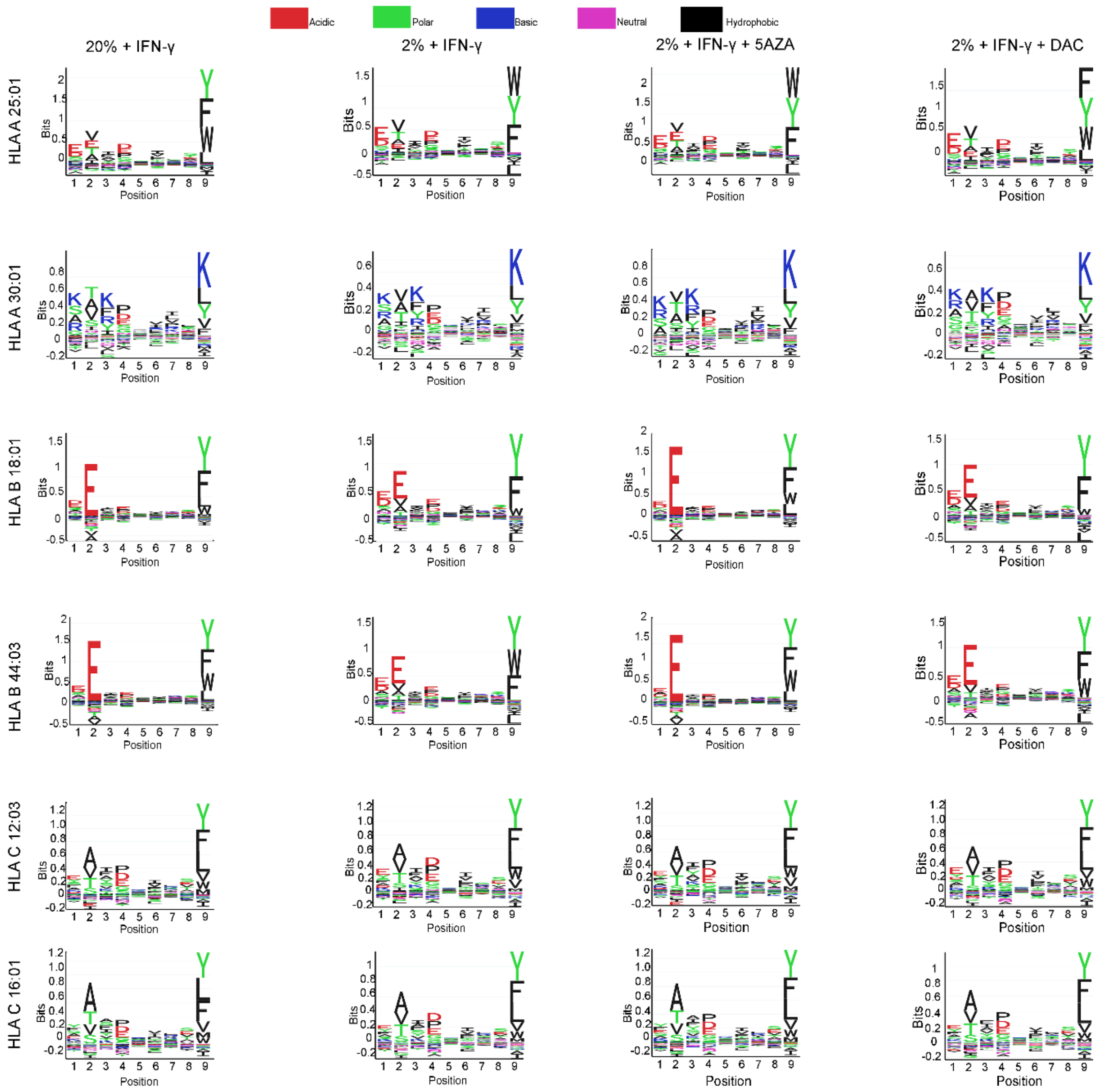


**Extended Data Figure 8: HLA allele-specific motif visualization following epigenetic pretreatment of hypoxic cells.**

Immunopeptidomics workflow and peptide identification were performed as described in Extended Data Fig. 2. A549 cells were cultured under indicated oxygen conditions with or without 5-AZA or DAC pretreatment for 48 h followed by 24 h IFN-γ stimulation. Gibbs clustering was used to generate allele-specific peptide-binding motifs. Sequence logos are arranged by HLA allele (rows) and condition (columns).

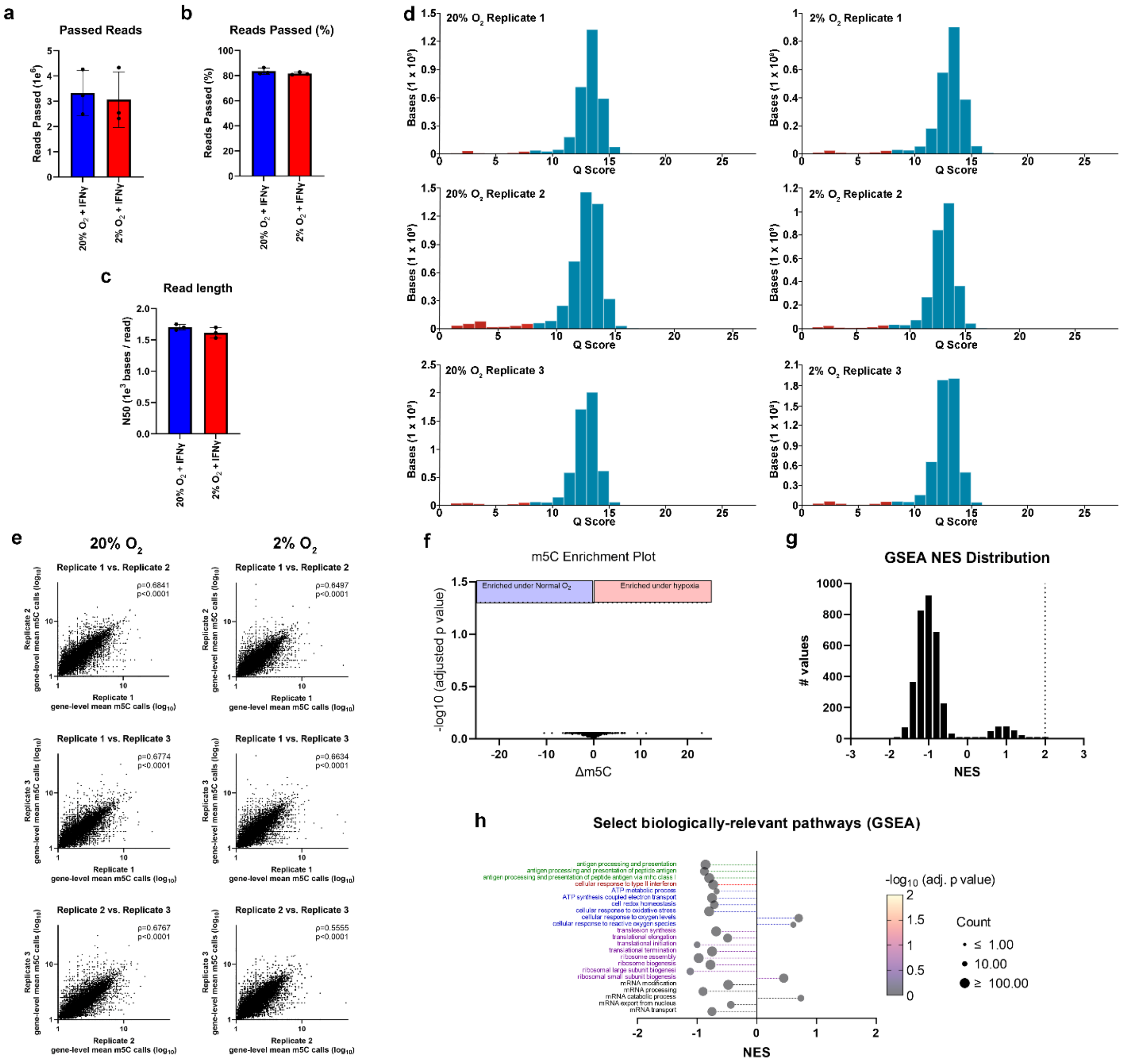


**Extended Data Figure 9: Nanopore direct RNA sequencing quality control and transcriptome-wide m5C analysis under normal $O_2$ and hypoxia**

**a-h,** A549 cells preconditioned for 48 h to 20% $O_2$ or 2% $O_2$ before treatment with IFN-γ for another 24 h were harvested, lysed, and RNA isolated for direct RNA nanopore sequencing with m5C-modified base calling. **a**, Total number of reads passing quality filters per sample following Dorado base-calling. **b**, Percentage of reads passing quality filters per sample. **c**, Median read length per sample following base-calling and alignment. **d**, Distribution of per-base Q scores across all reads for each biological replicate. **e**, Pairwise replicate concordance of gene-level mean m5C signal ($log_{10}$-transformed) within normoxic (left) and hypoxic (right) conditions. Each point represents one gene. Pearson correlation coefficients (ρ) and associated P values are shown. **f,** Transcriptome-wide differential m5C analysis displayed as Δm5C ($log_2$ fold change in mean methylation probability) versus $–log_{10}$ adjusted P value. Horizontal dashed line indicates the significance threshold after Benjamini–Hochberg false discovery rate (FDR) correction ($q < 0.05$). No transcripts meet this threshold. **g**, Distribution of normalized enrichment scores (NES) from gene set enrichment analysis (GSEA) performed on genes ranked by signed differential m5C signal. **h**, Selected biologically relevant pathways from GSEA, plotted by normalized enrichment score (NES). Dot size reflects gene set size and color indicates $–log_{10}$ adjusted P value. n=3 for all panels. For **a-c**, bars represent mean ± s.e.m

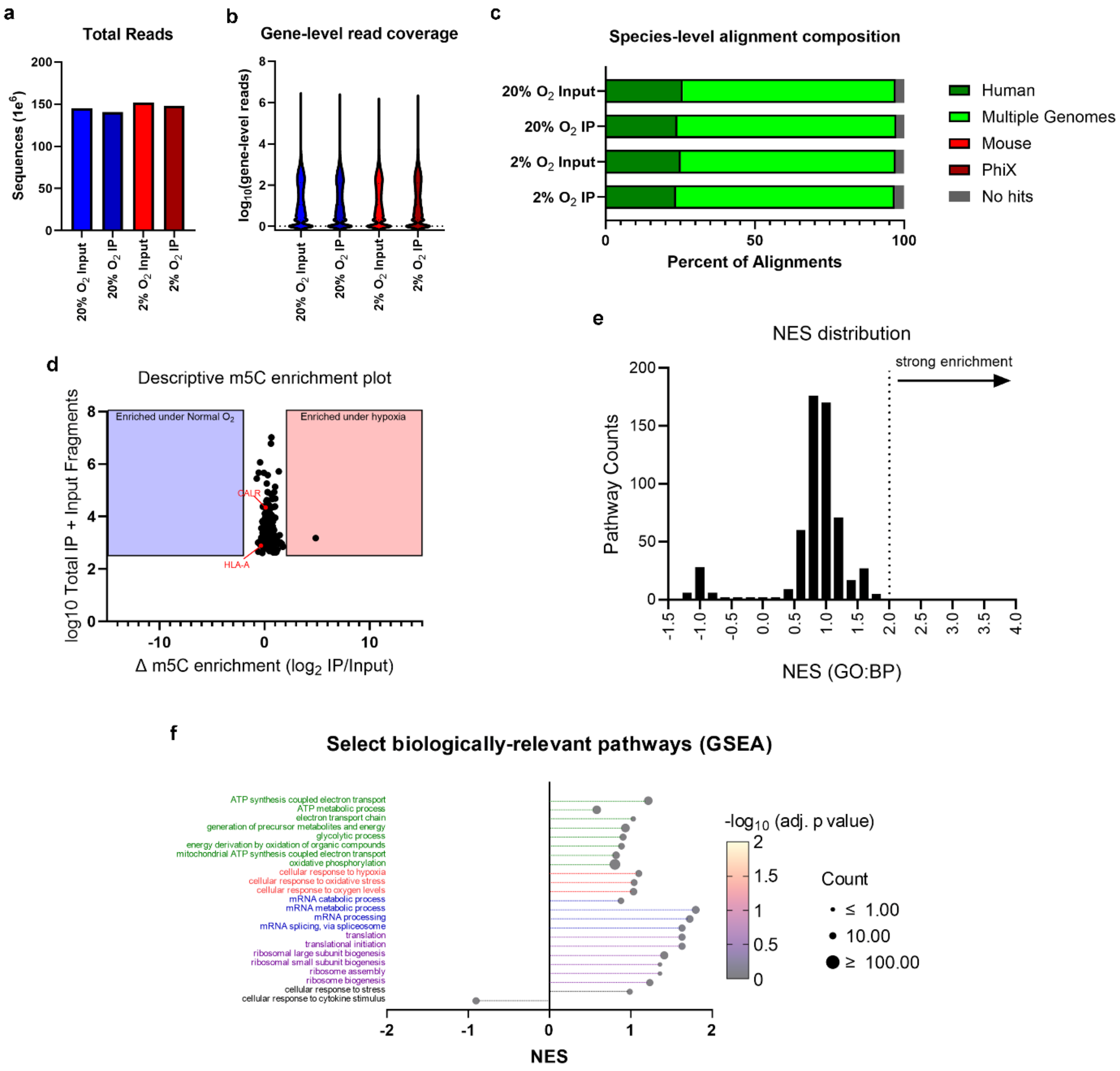


**Extended Data Figure 10: Quality control and transcriptome-wide m5C MeRIP-seq analysis under normal $O_2$ and hypoxia.**

A549 cells preconditioned for 48 h to 20% $O_2$ or 2% $O_2$ before treatment with IFN-γ for another 24 h were harvested, lysed, and RNA isolated for m5C MeRIP-seq analysis. **a**, Total sequencing depth per library (Input and IP) across 20% $O_2$ and hypoxic 2% $O_2$ conditions. **b**, Distribution of gene-level read coverage ($log_{10}$-transformed total reads per gene) for each library, demonstrating comparable transcriptome representation across conditions. **c**, Species-level alignment composition based on FastQ Screen analysis, showing the percentage of reads mapping to human, mouse, PhiX, multiple genomes, or remaining unmapped. **d**, Descriptive m5C enrichment plot showing Δm5C enrichment ($log_2$ IP/Input) versus total fragment abundance ($log_{10}$ IP + Input). Shaded regions denote transcripts relatively enriched under normoxia (left) or hypoxia (right). Representative transcripts are annotated. **e**, Distribution of normalized enrichment scores (NES) from gene set enrichment analysis (GSEA) performed on genes ranked by signed differential m5C signal. **f**, Selected biologically relevant pathways from GSEA, plotted by normalized enrichment score (NES). Dot size reflects gene set size and color indicates $-log_{10}$ adjusted P value.

### Supplementary Information

**Supplementary Table 1: Antibody reagents**

| *Target Protein* | *Clone Number* | *Company* | *Use* | *Catalog Number* |
|---|---|---|---|---|
| *HIF-1α* | D1S7W | Cell Signaling | W.B. | 36169S |
| *HIF-2α* | Polyclonal | R&D Systems | W.B. | AF2997 |
| *HIF-1β* | D28F3 | Cell Signaling | W.B. | 5537S |
| *FIH* | D19B3 | Cell Signaling | W.B. | 4426S |
| *PHD-2* | D31E11 | Cell Signaling | W.B. | 4835S |
| *α3* | D4Y90 | Cell Signaling | W.B. | 12446S |
| *β1* | E1K90 | Cell Signaling | W.B. | 13267S |
| *β2* | Polyclonal | ThermoFisher | W.B. | BS-9362R |
| *β5* | D1H6B | Cell Signaling | W.B. | 12919S |
| *β1i/LMP2* | E7J1L | Cell Signaling | W.B. | 87667S |
| *β2i/MECL-1* | EPR14902 | Abcam | W.B. | AB183506 |
| *β5i/LMP7* | D1K7X | Cell Signaling | W.B. | 13635S |
| *β-Actin* | 13E5 | Cell Signaling | W.B. | 4970S |
| *CALR* | D3E6 | Cell Signaling | W.B. | 12238 |
| *TAP1* | E4T4F | Cell Signaling | W.B. | 49671S |
| *Tapasin* | E6P2Z | Cell Signaling | W.B. | 66382S |
| *Erp57* | G117 | Cell Signaling | W.B. | 2881S |
| *ERAP1* | E6X1P | Cell Signaling | W.B. | 99979S |
| *IRF1* | D5E4 | Cell Signaling | W.B. | 8478S |
| *IRF2* | Polyclonal | Cell Signaling | W.B. | 4943T |
| *IRF3* | D6I4C | Cell Signaling | W.B. | 11904T |
| *IRF4* | D9P5H | Cell Signaling | W.B. | 15106T |
| *IRF5* | Polyclonal | Cell Signaling | W.B. | 20261T |
| *IRF6* | Polyclonal | Cell Signaling | W.B. | 6948T |
| *IRF7* | D8V1J | Cell Signaling | W.B. | 72073S |
| *IRF8* | D20D8 | Cell Signaling | W.B. | 5628T |
| *IRF9* | D2T8M | Cell Signaling | W.B. | 76684S |
| *IFNGR1* | E444 | Cell Signaling | W.B. | 10405S |
| *STAT1* | D1K9Y | Cell Signaling | W.B. | 14994T |
| *PSTAT1* | Y701 | Cell Signaling | W.B. | 7649S |
| *eIF2α* | D7D3 | Cell Signaling | W.B. | 5324S |
| *P-eIF2α (Ser51)* | D9G8 | Cell Signaling | W.B. | 3398T |
| *HLA-ABC (FITC)* | W6/32 | Biolegend | F.C. | 311404 |
| *HLA-ABC* | W6/32 | BioXCell | I.P. | BE0079 |
| *IFN-γ R α (PE)* | GIR-208 | Biolegend | F.C. | 308606 |
| *CA9* | M75 | ThermoFisher | I.F. | MA5-51609 |
| *G3BP1 (A549)* | EPR13986(B) | Abcam | I.F. | ab217225 |
| *Anti-rabbit IgG, Alexa Fluor 647* | Jackson ImmunoResearch | Polyclonal | I.F. | 711-607-003 |
| *Anti-mouse IgG, Alexa Fluor 594* | Jackson ImmunoResearch | Polyclonal | IF | 715-586-150 |
| *PSMB10/MECL1* | Abcam | EPR14902 | I.F. | ab183506 |
| *LMP7* | Abcam | EPR14482(B) | I.F. | ab180606 |
| *LMP2* | Abcam | EPR22042 | I.F. | ab242061 |
| *m5C* | Diagenode | 33D3 | I.P. | C15200081 |
| *m5C (FITC)* | Abcam | 5MC-CD | I.F. | ab179898 |
| *Anti-rabbit IgG, HRP-linked Antibody* | | Cell Signaling | W.B. | 7074S |

**Supplementary Table 2: Primer sequences used for quantitative PCR***

| GENE NAME | FORWARD | REVERSE |
|---|---|---|
| **PSMB5** | CCATACCTGCTAGGCACCAT | GCACCTCCTGAGTAGGCATC |
| **PSMB6** | CCTATTCACGACCGCATTTT | TCCCGGTAGGTAGCATCAAC |
| **PSMB7** | GCAACTGAAGGGATGGTTGT | AAACTAGGGCTGCACCAATG |
| **PSMB8** | GCTGCCTTCAACATAACATCA | CTGCCACCACCACCATTA |
| **PSMB9** | CTGGGACCAACGTGAAGG | ATGGCCAGAGCAATAGCG |
| **PSMB10** | GCTGCGGACACTGAGCTCA | TGGTTCCCAGGCACAAAGTGG |
| **TAP1** | GCAGTCAACTCCTGGACCACTA | CAAGGTTCCCACTGCTTACAGC |
| **TAPBP** | GAGCCTGTTCTCATCACCATGG | GTAGGCAAAGCTCAAGTCCAGC |
| **CALR** | TCAAGGAGCAGTTTCTGGACGG | CGATCCTGGCTTGTCTGCAAAC |
| **B2M** | CCACTGAAAAAGATGAGTATGCCT | CCAATCCAAATGCGGCATCTTCA |
| **PDIA3** | GTCAGCCACTTGAAGAAGCAGG | TAGGAACTCGGAGTGAGCCTCA |
| **IRF1** | GAGGAGGTGAAAGACCAGAGCA | TAGCATCTCGGCTGGACTTCGA |
| **IRF7** | CCACGCTATACCATCTACCTGG | GCTGCTATCCAGGGAAGACACA |
| **IRF8** | AGGTCTTCGACACCAGCCAGTT | GCACGAGAATGAGTTTGGAGCG |
| **IRF9** | CCACCGAAGTTCCAGGTAACAC | AGTCTGCTCCAGCAAGTATCGG |
| **ACTB** | CACCATTGGCAATGAGCGGTTC | AGGTCTTTGCGGATGTCCACGT |

* All primers were designed to amplify human transcripts and were validated by melt-curve analysis. ACTB was used as the endogenous control for ΔΔCt normalization.

**Supplementary Table 3: CRISPR/Cas9 gRNA sequences***

| *Target* | *Gene* | *crRNA sequence (5' → 3')* |
|---|---|---|
| *HIF-1α* | **HIF1A** | CCUCACACGCAAAUAGCUGA |
| *HIF-1α* | **HIF1A** | ACAGUAACCAACCUCAGUGU |
| *HIF-2α* | **EPAS1** | CAAGGCCUCCAUCAUGCGAC |
| *HIF-2α* | **EPAS1** | GUUGACAGUACGGCCUCUGU |

*All guide RNAs were synthesized by Integrated DNA Technologies (IDT) as Alt-R CRISPR-Cas9 crRNAs and used in ribonucleoprotein (RNP) complexes with Alt-R tracrRNA and S.p. Cas9 nuclease V3. For both HIF-1α and HIF-2α targeting, two independent guides were used simultaneously to increase editing efficiency. The HIF-1α / HIF-2α double KO line was generated sequentially by targeting EPAS1 in a validated HIF1A knockout clone.

**Supplementary Table 4: Custom Steallaris™ FISH Probes**

| | *PSMB8* | *PSMB9* | *PSMB10* |
|---|---|---|---|
| *Probe 1* | ctctcggaggaggaagtgaa | tgggacagcgcacaacgc | tctgctaggcttcacgtc |
| *Probe 2* | atacatctagtagcgccatg | tggggcactggtttccaa | ggccagtgagcagctaaa |
| *Probe 3* | gagaaactgtagtgtcctgg | agcatccctgcaaggcac | tgagtcggccagacaagc |
| *Probe 4* | gtaaagcgagctctggagat | gtaagtccccggttggtg | ttctcgaaggagaagccc |
| *Probe 5* | ccagggactggaagaattct | gtcaaactccactgccat | ggaggacgcgttccaatg |
| *Probe 6* | tctcaatctgaacgttcctt | agaacccatcacaacgcc | tgcgtgagggaccttgag |
| *Probe 7* | ctccatgctggaacttgaag | cctgcagacactcgggaa | tatcggcgcccagaatga |
| *Probe 8* | ccgtaaggcactaatgtagg | aacactcggttcaccacc | ccgaatcgttagtggctc |
| *Probe 9* | gcaggtaagggttaatctca | tgcagcggggacagcttg | cttctcgcagctcttgtc |
| *Probe 10* | cagtactgacagtctgctgc | agtgcacagtagatgcgc | ttgggggcgatgaagtgg |
| *Probe 11* | tccatttcgcagatagtaca | catcagcagctgaaccag | catccgtgtggtcatctc |
| *Probe 12* | ctgccgacactgaaatacgt | catgtcggccacggcttg | gcccgtagataacgcgtg |
| *Probe 13* | catgttcatccacgtagtag | atggagctccagctggta | gatgcgagtgaccgtggc |
| *Probe 14* | gaacatatttcctgagagcc | gcagcagccaaaacaagt | ctgaagagcgtctggcgc |
| *Probe 15* | aggcataagtgttcccacta | gctgatatttctcaccac | gatcagcgatgcacccac |
| *Probe 16* | taagattaggccgatagcca | gcagacaagtcctctcga | cagtcaggtctacgccgc |
| *Probe 17* | aaggtcataggcctcttcag | agccagctaccatgagat | cacaccgtagagctgcgg |
| *Probe 18* | tctgtgagtggcataagcaa | ctgacctccttcacgttg | ctgtaggagccatgggga |
| *Probe 19* | tactttcacccaaccatctt | tcccagggttccatatac | agggctgtgaagggcaga |
| *Probe 20* | caggtcactgacatctgtac | aaggctgtcgagtcagca | gtcttctagcaccgccag |
| *Probe 21* | caagacctcccagaggagac | gataaaggtgctgccgga | cgtcatgttcggctggaa |
| *Probe 22* | cttggacttaacgtggctta | gctgcatccacataacca | caagatcccggcggtgac |
| *Probe 23* | tactctctctttggctcagg | gagacatgcctggcttat | cacatgcgtccacattgc |
| *Probe 24* | acacgcagaagatgcacttc | tctgtggtgaagcgcctg | ttggcgccagtctttgtg |
| *Probe 25* | tgggggaaatgcttgttcaa | catggccagagcaatagc | tgagctcagtgtccgcag |
| *Probe 26* | ttagtggggcacccagaaac | caggtagatgacaccccc | ctcttcacgggctctgtg |
| *Probe 27* | | ccggcagctgtaatagtg | caaagtggtagcggccag |
| *Probe 28* | | agatgactcgatggtcca | acagtttcctccactagc |
| *Probe 29* | | ttttggcagttcattgcc | tttctgggtttattcccc |
| *Probe 30* | | tttggccctagagagttt | |

* https://oligos.biosearchtech.com/products/rna-fish/stellaris-probe-designer
